# Scalable construction of near-wild-type brain atlases enables variability-aware neuronal identification in *C. elegans*

**DOI:** 10.64898/2026.11.12.766186

**Authors:** Zikai Yu, Wenjing Yang, Yesol Moon, Jingting Liang, Sihoon Moon, Zhenning Yang, Hyun Jee Lee, Lizeth Parra Arroyo, Minghai Ge, W. Alexander Calhoun, Shiv Kumar, Yun Zhang, Hang Lu

## Abstract

Matching cells across individuals is essential for comparing cellular data but becomes challenging when cell positions vary between animals. Here we introduce a framework for constructing probabilistic neuronal atlases by integrating partially overlapping observations from sparsely labeled, near-wild-type strains to reconstruct cellular organization from incomplete observations. Using five strains imaged without compression in a microfluidic array, we built an atlas of 129 head neurons (55 classes) that captures positional variability and infers spatial relationships even between neurons never observed together. We find that variability is spatially structured with neurons in compact regions being more stereotyped, while neuronal relative positions are more stable than their absolute positions. Exploiting these statistical relationships, variability-weighted identification improved top-1 accuracy to 86.3%, approaching human inter-annotator agreement, with the correct identity among the top three candidates in 98.2% of cases. Relationships stitched across strains reproduced observations in a held-out strain, and the atlas supplied correct identities in males and in pan-neuronal calcium imaging in vast majority of cases. Our extensible framework enables community-built atlases and offers a strategy for cell identification when anatomical variability limits template-based registration.

## Introduction

Comparing cellular data across individuals, experimental modalities, and laboratories requires a common reference frame in which the data can be aligned and corresponding cells identified. This challenge arises in many applications, including whole-brain functional imaging, connectomics, and spatial omics. In several model organisms, substantial efforts have established standardized anatomical references, including the Allen Mouse Brain Common Coordinate Framework [1], Z-Brain and mapZebrain in zebrafish [2, 3], and the JRC2018 templates and the FlyWire connectome in *Drosophila* [4, 5]. Despite differences in anatomical scale, spatial resolution, and data modality, these approaches largely establish correspondence among samples and datasets by registration to a reference template, placing biological information in a common frame for understanding how these brains develop and function. For such atlases to be broadly useful, they must satisfy three criteria. The first is fidelity, which cannot be represented by a single idealized arrangement. The others are efficiency and accessibility, which require that other researchers be able to use the atlas, extend it to new conditions, or build comparable atlases of their own without specialized instrumentation, large number of transgenic crosses, or extensive recalibration. Existing tools implicitly assume a trade-off among these criteria. We argue that this apparent trade-off arises from particular design choices in existing atlas-building tools.

Approaches to neuronal identification and atlas construction rely on two complementary strategies: spatial registration and molecular labeling. Spatial registration establishes correspondence by aligning anatomical features to a common reference, such as a template or a standard coordinate system. This strategy has been used to build brain atlases across model organisms, including whole-brain atlases of the nematode *Caenorhabditis elegans* (*C. elegans*) [6–9]. For example, Toyoshima et al. [9] identified head neurons in 311 adult worms using 35 cell-specific promoters and combined their positions into a comprehensive neuronal atlas. Although registration-based approaches can integrate data across samples, their dependence on a common reference can introduce errors and distortions arising from anatomical variability, sample deformation, and differences in imaging conditions. Molecular labeling approaches instead assign neuronal identity through combinations of fluorescent markers, as in Brainbow [10] and, in *C. elegans*, NeuroPAL [11], or through transcriptomic profiles, as in spatial omics [12]. These approaches provide reliable identity information, but they require specialized transgenic labeling and imaging protocols that can perturb physiology and limit accessibility. High-resolution methods such as electron-microscopy reconstruction [13–15] and expansion microscopy [16, 17] provide complementary anatomical information but remain difficult to scale to large and live populations. These strategies therefore anchor identity to a canonical reference, but involve trade-offs. For instance, registration is sensitive to anatomical variability and experimental conditions, and extensive labeling limits applicability and accessibility. Neither exploits the relative arrangement of neurons, which is more conserved across individuals.

Testing whether relational information can support atlas construction requires a system in which ground-truth correspondence exists. *C. elegans* is an ideal test bed: the 302 neurons of the hermaphrodite are individually identifiable by anatomical position [13] and gene expression [18], their connectivity is fully mapped [13, 14, 19], and the cell lineage is fully characterized [20, 21]. However, neuronal positions vary substantially between individuals [9, 11, 22, 23], so the reliability of registration-based methods depends strongly on how variable each cell is. The animal’s optical transparency enables fluorescence imaging without sectioning, its short, stereotyped life cycle yields synchronized populations that can be imaged at scale, and its genetic toolkit allows precise labeling of individual neurons or neuronal classes. These features have enabled studies that integrate spatial, molecular, morphological, and functional information [18, 24, 25], including the interpretation of whole-brain calcium imaging [26–28]. In all of these cases, interpretation depends on accurate neuronal identification and therefore on the quality of the atlas used.

Here we describe a pipeline that satisfies fidelity, efficiency, and accessibility together by anchoring the atlas to relative spatial information instead of absolute positions alone (**Fig. 1**). We combine sparsely labeled near-wild-type strains, high-throughput imaging, BFP-verified consensus annotation, and label-free anatomical feature-anchored alignment into a graph atlas that can be continuously improved or extended. Building on this framework, we generated a head atlas for young-adult hermaphrodite as an example. From five strains, the atlas covers 129 head neurons in 55 classes. We show that relationship stitched across strains, including those between neurons never observed together, is as good as that produced from a held-out strain. The atlas also contains positional variability, which we show is highly structured and influenced by where the neurons are located and how dense the cells are locally. Weighting the identification of neurons by this variability greatly improves the accuracy of the identity prediction. We further demonstrate that a hermaphrodite-derived atlas reliably identifies neurons in males, and that automatic, accurate neuron identification in whole-brain functional imaging experiments can substantially reduce the expertise and labor that neuron identification requires.

**Fig. 1:**
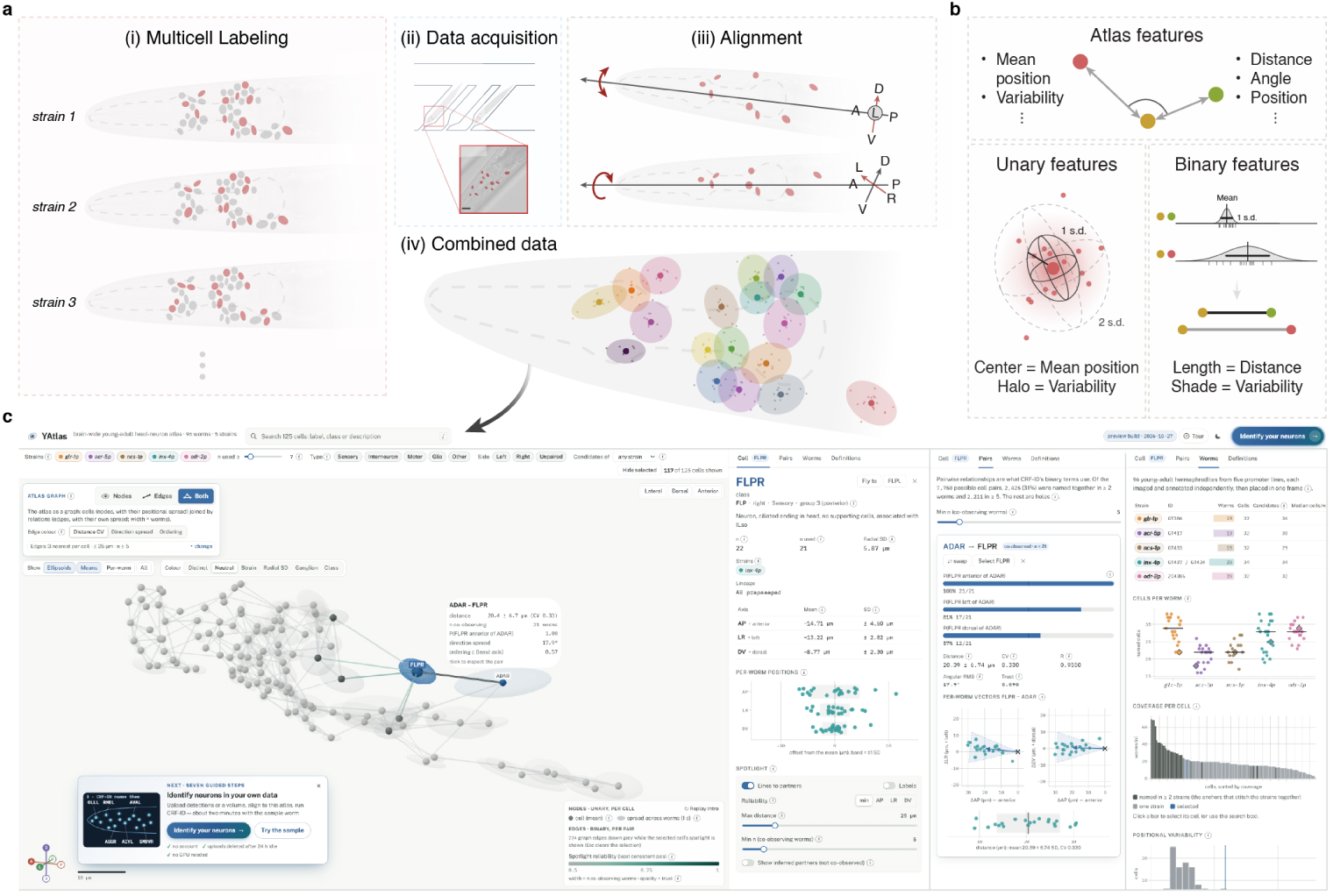
A modular pipeline for building probabilistic brain-wide atlases from partially overlapping multi-cell strains. **a.** To address the challenges of fidelity, efficiency, and accessibility, our method utilizes three main approaches. (i) Five multi-cell strains are used to cover the brain-wide neural distribution. Each strain is labeled with a different subset of neurons, where they share a number of neurons with at least one other strain for downstream data alignment. (ii) Then, the worms are imaged using a microfluidic device designed to allow high-throughput data acquisition with optimal orientation and minimal deformation. (iii) Lastly, the obtained data are aligned into the same physical space using a two-step alignment: registration of data within each strains, followed by cross-strain alignment. The resulting combined data has information about both the mean position of neurons (center points shown in the figure), as well as the positional variability (depicted as ellipsoids in the figure). **b.** The brain-wide atlas provides anatomical information at both the unary level (features specific to each neuron) and the binary level (features between neuron pairs). More importantly, the data-driven nature of the atlas-building pipeline allows us to estimate the positional variability of covered neurons (shown as halo shade around each neuron) and how much the relative position changes from individual to individual (indicated by the shade of edges in the graph). **c.** The brain-wide atlas is a graph-like structure, in which neurons are nodes and pairwise positional relationships are edges. The atlas stores unary features at nodes (mean position and its variability) and binary features at edges (pairwise distance and direction, each with its variability). The atlas is provided with a GUI, where each node is represented at its mean position (the center point), with the positional variability shown by the halo radius. The distances between these nodes and their variability are represented by the distance and the color of the edge respectively, with higher correlation (lower variability) shown as a darker color. A new dataset takes the same form with unlabeled nodes, so the identification becomes a graph-matching problem.

## Results

### A modular pipeline for building probabilistic brain-wide atlases from multi-cell data

To address the challenges of fidelity, efficiency, and accessibility that existing atlas-building approaches struggle to balance, we developed a modular pipeline that produces probabilistic atlases of the head neurons and can be extended without being rebuilt (**Fig. 1**; Methods). This platform includes a set of multi-cell reporter strains (**Fig. 1a (i)**), a high-throughput imaging device (**Fig. 1a (ii)**), a structured protocol for establishing ground-truth via human annotators, an anatomy-anchored alignment procedure (**Fig. 1a (iii)**), unary and binary level features extracted from the combined data (**Fig. 1a (iv) and b**), and a graphical representation (**Fig. 1c**) built on relative positional constraints. Each of these tools is designed to address the challenges brought by specific sources of distortion from the idealized atlases in each sample (**Supplementary Fig. 1**).

A single pan-neuronal marker makes neighboring neurons difficult to distinguish during annotation, while a multicolor code imposes a transgenic load that moves the animals measurably away from wild type. Therefore, we distribute coverage of the head across several multi-cell strains, each carrying one nuclear-localized reporter, which keeps annotation tractable at minimal genetic cost and requires only about ten animals per strain for a stable atlas (**Supplementary Fig. 2**). Requiring fewer animals provides flexibility in the imaging setup, but agar-pad mounting compresses each animal differently, and this variation becomes indistinguishable from biological variability once the data are pooled. We therefore imaged animals in an array-style microfluidic device that holds every worm in the same uncompressed geometry and admits more than 100 animals per five-minute loading, so that data collected on different days and from different strains are physically comparable before any computation (**Supplementary Figs. 3 and 4**). On the data processing side, annotators working without shared criteria may apply different features and arrive at confidently different labels. To prevent such discrepancy, we detected nuclei automatically using a widely used algorithm (StarDist, **Supplementary Fig. 5**), and established per-strain annotation criteria in advance using auxiliary BFP lines and that do not contribute in building the atlas itself [29]. Finally, we had at least three annotators label every volume independently before reconciling their assignments into probabilistic labels, that retain both confidence and the disagreement between annotators (**Supplementary Figs. 6 and 7**). When consolidating the curated data, two multi-cell strains may share only a handful of labeled neurons, too few for registration to constrain a transform, and the resulting errors can inflate the variability reported by the atlas. Thus, we anchored alignment to the pharyngeal bulbs and the worm centerline, features every animal possesses regardless of its expression pattern, which leaves a single rotational degree of freedom per animal and yields an atlas independent of the order in which data are combined (**Supplementary Figs. 8–11**). The aligned data are stored as a graph in which each neuron is a node carrying its mean position and variability, and each pair of neurons is an edge carrying the distance and direction between them with their variability across animals (**Fig. 1b and c**). With this approach, identifying neurons in a new dataset becomes a graph-matching problem and extending the atlas becomes graph merging. Finally, the brain-wide atlas is provided in a graphical user interface (GUI) (**Fig. 1c**), allowing any user to access the related information and compare it with their data.

### Multi-cell strains preserve wild-type physiology and behavior

Our current brain-wide atlas is built from five strains, driven by the *glr-1*, *acr-5*, *inx-4* (with *mbr-1*), *ncs-1*, and *odr-2*promoters (**Fig. 2a**); a sixth, *flp-3* (with *flp-7*, *sro-1*, and *nmr-1*), was held out for validation. Each strain labels an average of 30 head neurons drawn from major ganglia, and since every promoter had been characterized previously [18, 30–32], the set of cells expected in each strain was known before annotating them (**Supplementary Fig. 12 and 13**). Together, the five atlas-building strains cover approximately 66 % of head neuron classes (55 classes, 129 cells); further coverage depends on the availability of suitable promoters and is not inherently limited by the pipeline. The coverage map shown in **Fig. 2b** represents what the atlas can measure directly and what it has to infer. Each strain shares a number of neurons with at least one other strain, which allows data from different strains to be brought into a common frame for alignment. However, the overlaps are small (as few as six neurons), and most neurons only appear in one strain, so the majority of neuron pairs are never observed together. The positional relationships between these pairs are inferred by aligning strains during data processing. As shown in the next section, these inferred relationships are as accurate as those measured directly within individual animals.

**Fig. 2:**
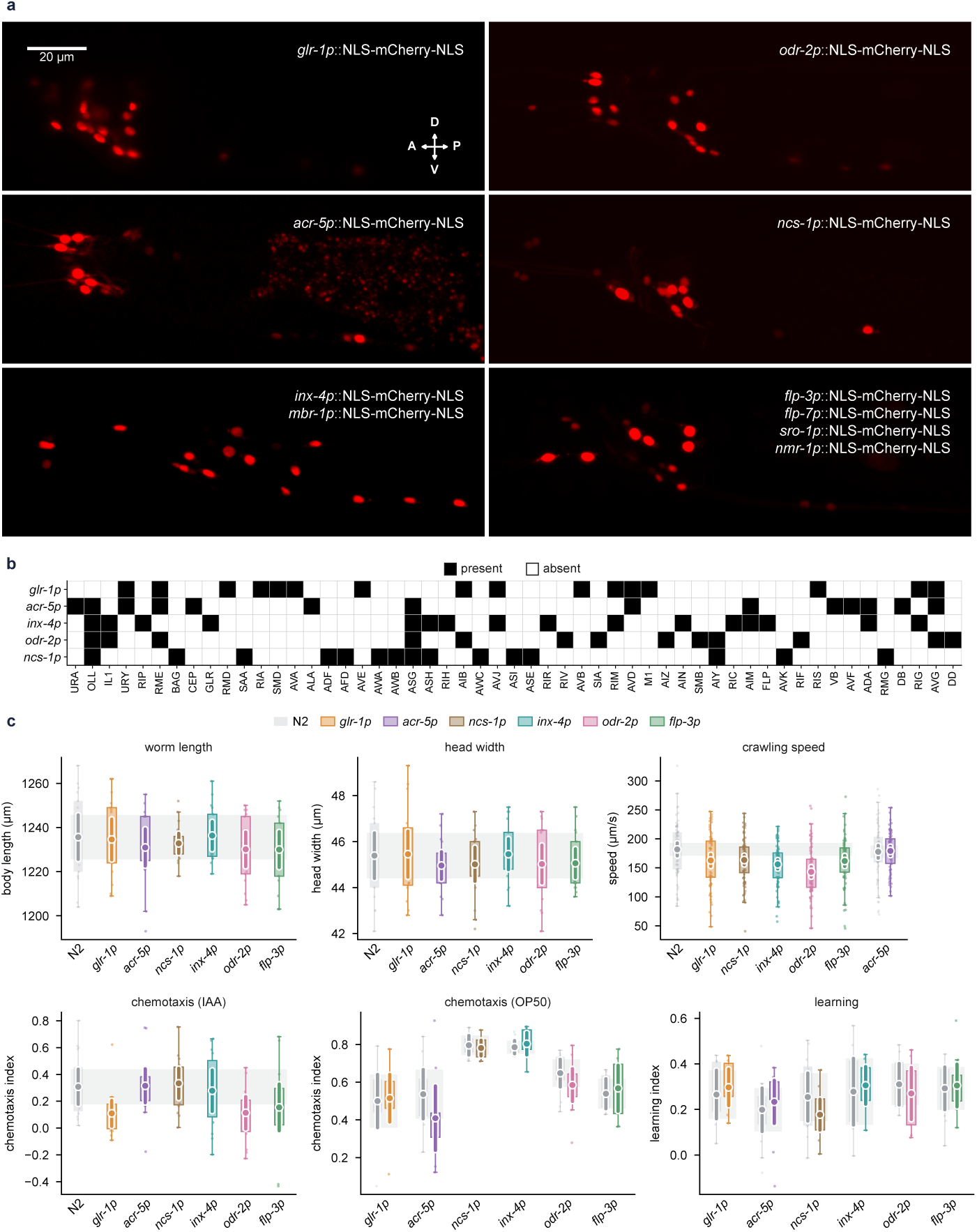
Multi-cell strains preserve wild-type physiology and behavior. **a**. Representative images (maximum intensity projections) of the five strains used for atlas construction (*glr-1p*, *odr-2p*, *acr-5p*, *ncs-1p*, and *inx-4p & mbr-1p*) and the heldout strain used for validation (bottom right, *flp-3p & flp-7p & sro-1p & nmr-1p*). **b**. Coverage map: matrix rows correspond to different atlas construction strains, columns correspond to neurons (cell classes) labeled by fluorescent reporter (black: present, white: absent). Neuron classes are ordered from anterior to posterior. **c**. Reporter strains compared with wild type (N2); strains are named by their first promoter (**a**). Top, body length, head width and crawling speed on food. Bottom, chemotaxis toward isoamyl alcohol (IAA; navigation index of single worms on agar plates), chemotaxis toward the odor of *E. coli* OP50 (microdroplet assay) and aversive olfactory learning (learning index: OP50-versus-PA14 choice index of naive minus PA14-trained worms; microdroplet assay). Boxes, interquartile range; whiskers, 1.5x IQR; dots, individual worms (body length, head width, crawling speed, IAA) or replicate assays (OP50, learning); circles and thick bars, mean and 95% confidence interval of the mean. Gray bands, 95% confidence interval of the N2 control, drawn across the strains tested with it. The OP50 chemotaxis assays and learning assays have paired N2 control for every strain. *n* per group: body length and head width, 20 worms; crawling speed, 76-84 worms per strain and around 70 N2 worms; IAA, around 15 worms per strain; OP50, 9-12 replicates per strain across 3-4 days; learning, 9-12 replicates per strain. More results are reported in **Supplementary Fig. 14**.

Because the atlas is intended to describe wild-type anatomy, the strains that we used to build the atlas should be close to N2 reference. To confirm this, we measured body length and head width, the dimensions most relevant to where head neurons sit. None of the strains differed significantly from N2 (**Fig. 2c**). In a separate locomotion assay on food, we measured peristaltic speed, defined as the absolute peristaltic track length divided by the elapsed time, and again found no significant difference from N2. We also assessed higher-level functions. In both chemotaxis assays toward isoamyl alcohol (IAA) and the standard food source *E. coli* OP50, every strain performed comparably to N2. Finally, we tested the learning capabilities of each strain, which also showed no significant difference from that of N2. Additional measures, including brood size and extra associative learning results, are reported in **Supplementary Fig. 14**. Taken together, these results indicate that the single-reporter strains used here are physiologically and behaviorally close to wild type, suggesting that the anatomy extracted from them is representative of N2 and not substantially altered by the transgenic modification.

### Cross-strain integration recovers valid positional relationships for neuron pairs that are never jointly observed

We first asked whether a data-driven atlas is necessary at all, or whether an existing positional reference would serve as well. For each construction strain, every worm was held out in turn and its neurons were identified against three references: a static atlas built from OpenWorm coordinates [33], an atlas derived from NeuroPAL [11] (truncated to the labeled subset of head neurons), and an atlas built from the remaining worms of the same strain (**Fig. 3a**). The data-driven atlas outperformed both alternatives in every strain, and by a wide margin: correspondence with manual identities was centered near 0.85 to 0.90 for the data-driven atlas, against roughly 0.4 to 0.6 for NeuroPAL and 0.3 to 0.4 for OpenWorm. The same test also establishes that each strain is internally consistent, since an atlas built from a subset of the data predicts the rest with high accuracy. When we considered the three highest-ranked candidates, correspondence reached 0.95 or above in every strain, and 0.98 or above at rank five (**Fig. 3b and Supplementary Fig. 15–16**). This shows that even if the single best assignment is wrong, the correct identity is almost always within a short list of which a user can review.

**Fig. 3:**
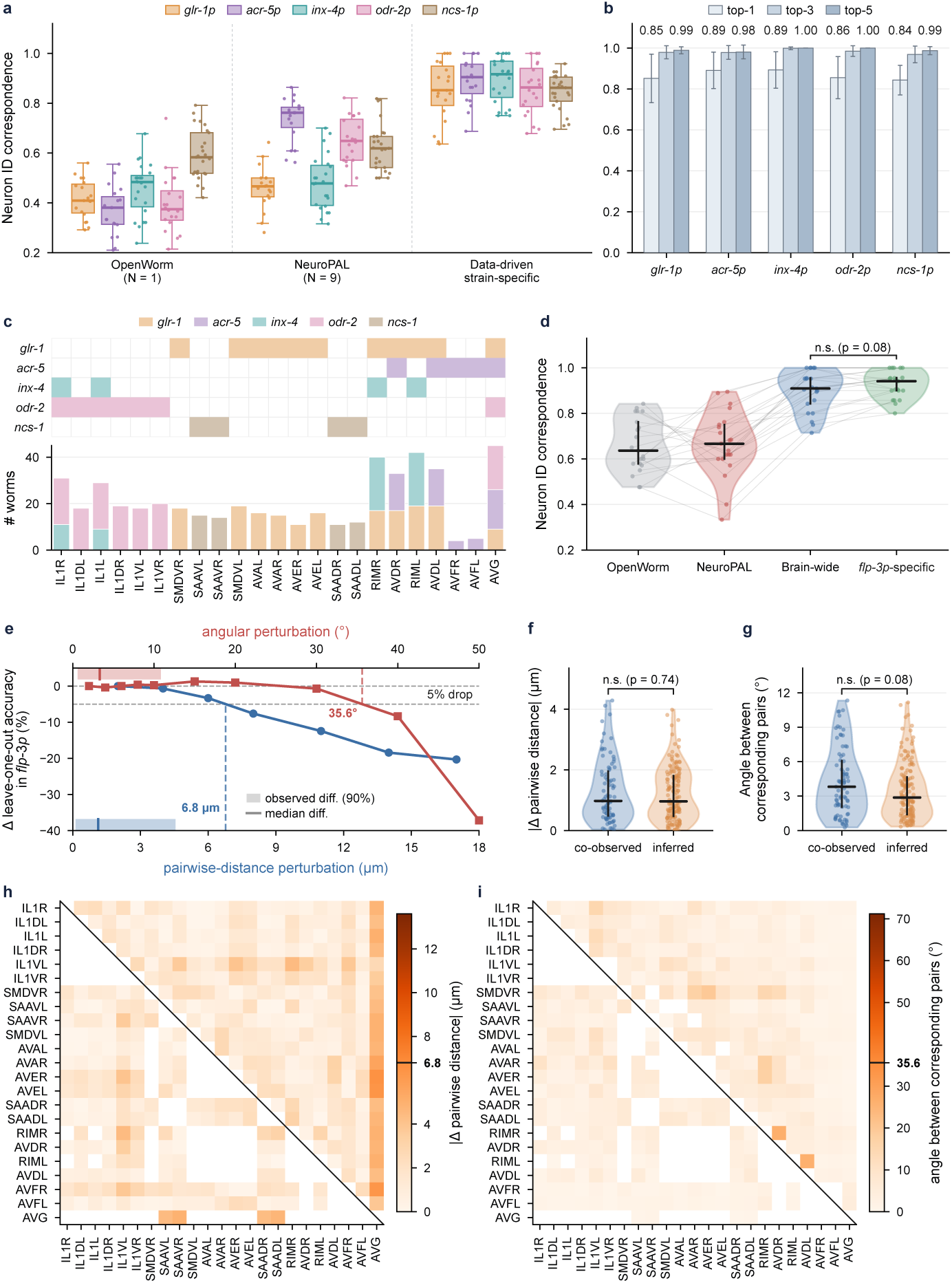
Cross-strain integration recovers valid positional relationships for neuron pairs never jointly observed. **a**. Strain-specific neuron identification accuracy comparing three sources of reference data: OpenWorm, NeuroPAL, and the data-driven atlas (this work). OpenWorm and NeuroPAL atlases were truncated to only the neurons labeled by each reporter strain. For obtaining data-driven identification accuracy, data for each worm to be tested was excluded from the atlas when inferencing that specific worm (leave-one-out validation). Box plots show median and interquartile range; whiskers, 1.5x IQR; points, individual worms; n = 21-25 per strain. **b**. Top-1 (neuron ID correspondence), top-3, and top-5 identification accuracy for the data-driven atlas. Bars, mean; error bars, SEM; values above bars are means. **c**. (Top) The majority of neurons (columns) expressed in the held-out validation strain (*flp-3p & flp-7p & sro-1p & nmr-1p*) is present in at least one atlas construction strain (rows). The four SIB neurons expressed in the validation strain are not covered by any atlas strain and were excluded. (Bottom) Number of worms in which each *flp-3* neuron was observed, stacked by contributing strain. **d**. Identification accuracy on *flp-3* neurons using the OpenWorm atlas, the NeuroPAL atlas, the brain-wide atlas built from the five construction strains without *flp-3p* data, and the *flp-3* -specific atlases built by leave-one-out validation. The analysis is limited to the candidate list of this validation strain. The brain-wide atlas performs comparably to the *flp-3p*-specific atlas. Violins, distribution; horizontal bars, median; gray lines connect the same worm across atlases; n.s., two-sided Wilcoxon signed-rank test, P = 0.08, N = 23. **e**. *flp-3* leave-one-out accuracy as a function of controlled perturbation applied to pairwise distances (blue) and to pairwise angles (red) using the *flp-3* -specific atlas. Shaded region, 90th percentile of the observed difference between the brain-wide atlas and *flp-3* -specific atlas; dotted line, median observed difference; dashed line, perturbation producing a 5 % accuracy drop (6.8 *µ*m; 35.6*^◦^*). **f** and **g**. Absolute difference in pairwise distance (left) and angle between corresponding pair vectors (right) between the brain-wide and *flp-3* -specific atlases, for pairs co-observed in at least one construction strain versus pairs whose relationship was inferred through alignment alone. n.s., two-sided Mann–Whitney U test; P = 0.74 (distance), P = 0.08 (angle); N = 80 for co-observed pairs, N = 155 for inferred pairs. **h** and **i**. Residual matrices for pairwise distance (**h**) and angles across neuron pairs (**i**). Upper triangle, all neurons pairs considered; lower triangle, pairs never co-observed within a single construction strain; color scale, residual magnitude, with the tick at the 5 % threshold from **f**.

One remaining question arose from a design risk introduced by using multi-cell strains: because each strain labels only part of the head, many neuron pairs are never observed together in any animal, and their relationship in the combined atlas exists only because the strains were placed in a common coordinate system. To test whether those relationships are accurate, we imaged a sixth strain, *flp-3* (with *flp-7*, *sro-1*, and *nmr-1*), that was not used in construction, and asked whether an atlas built from the other five could identify its neurons. *flp-3* suits this purpose because most of its labeled neurons are already covered by the brain-wide atlas, and the coverage is spread across all five construction strains **(Fig.3c**). Each covered neuron is supported by observations from between roughly ten and forty animals, and the four SIB neurons that *flp-3* labels but are not covered by any other strain, were excluded.

Identification accuracy on *flp-3* neurons was indistinguishable between the brain-wide atlas which had never seen a *flp-3* animal, and a strain-specific atlas built from *flp-3* by leave-one-out (**Fig. 3d**; P = 0.08). Additionally, both showed an accuracy significantly higher than the OpenWorm and NeuroPAL references. The combined atlas therefore can recover the structure of a strain it was never trained on, including neuron pairs it could only have obtained by stitching. To interpret how close the brain-wide atlas and the *flp-3* specific atlas are, we needed to know how much a positional relationship can change before the identification results lose accuracy. We perturbed the pairwise distances and pairwise angles in the *flp-3* -specific atlas by a controlled amount and measured the effect on a leave-one-out accuracy test (**Fig. 3e**). The accuracy did not show a significant decrease until distances were shifted by several micrometers, and fell by five percentage points at only 6.8 *µ*m; for angles, the corresponding threshold was 35.6*^◦^*. The observed differences between the brain-wide and *flp-3* -specific atlases showed medians near 1 *µ*m and 3*^◦^*, which sit well inside those tolerances.

Finally, we compared the relationships between neurons, separating pairs that were co-observed in at least one construction strain from pairs whose relationship was inferred through alignment alone. The two groups did not differ in distance residual (P = 0.74) or in angular residual (P = 0.08) (**Fig. 3f and g**), and the full residual matrices show no structure that would single out the inferred pairs (**Fig. 3h–i and Supplementary Fig. 17**). These results show that the relationships that the brain-wide atlas assemble across strains are as accurate as the ones measured directly. This is the property that makes the multi-cell approach practical, allowing the atlas to capture relationships between neurons without requiring users to observe every pair in the same animal or image the same strains used for atlas construction.

### Quantified positional variability confirms the robustness of relational features and improves automated neuron identification

Now with the brain-wide atlas validated, we examined what it reports. For every neuron represented in the atlas, aligned positions across animals are used to construct a three-dimensional distribution, shown as an ellipsoid in the atlas coordinate system (**Fig. 4a**). Larger ellipsoids indicate greater variability. For many neurons, the expected position distribution overlaps with that of their nearest neighbors. Variability is greatest in the retrovesicular ganglion, where neurons are sparse, and smallest in the anterior ganglion, where they are densely packed. Along the anterior-posterior axis, variability rises where local neuron density falls (**Fig. 4b and Supplementary Fig. 18**). Across all covered neurons, variability and local neuron density are negatively correlated (**Fig. 4c**; *r* = −0.57). Neurons packed among many neighbors have less room to move. This is the kind of measurement that an atlas built from a few animals cannot make, and it agrees with the substantial individual variability documented in other whole-brain datasets [11, 23]. The same variability, together with expression mosaicism [35] and local density, accounts for much of the disagreement between human annotators on the same image (**Supplementary Fig. 19**). This visualization of variability not only illustrates the positions of neurons, but also the overall challenge of annotation based on position alone and specific regions where variability makes relying on position alone unreliable.

**Fig. 4:**
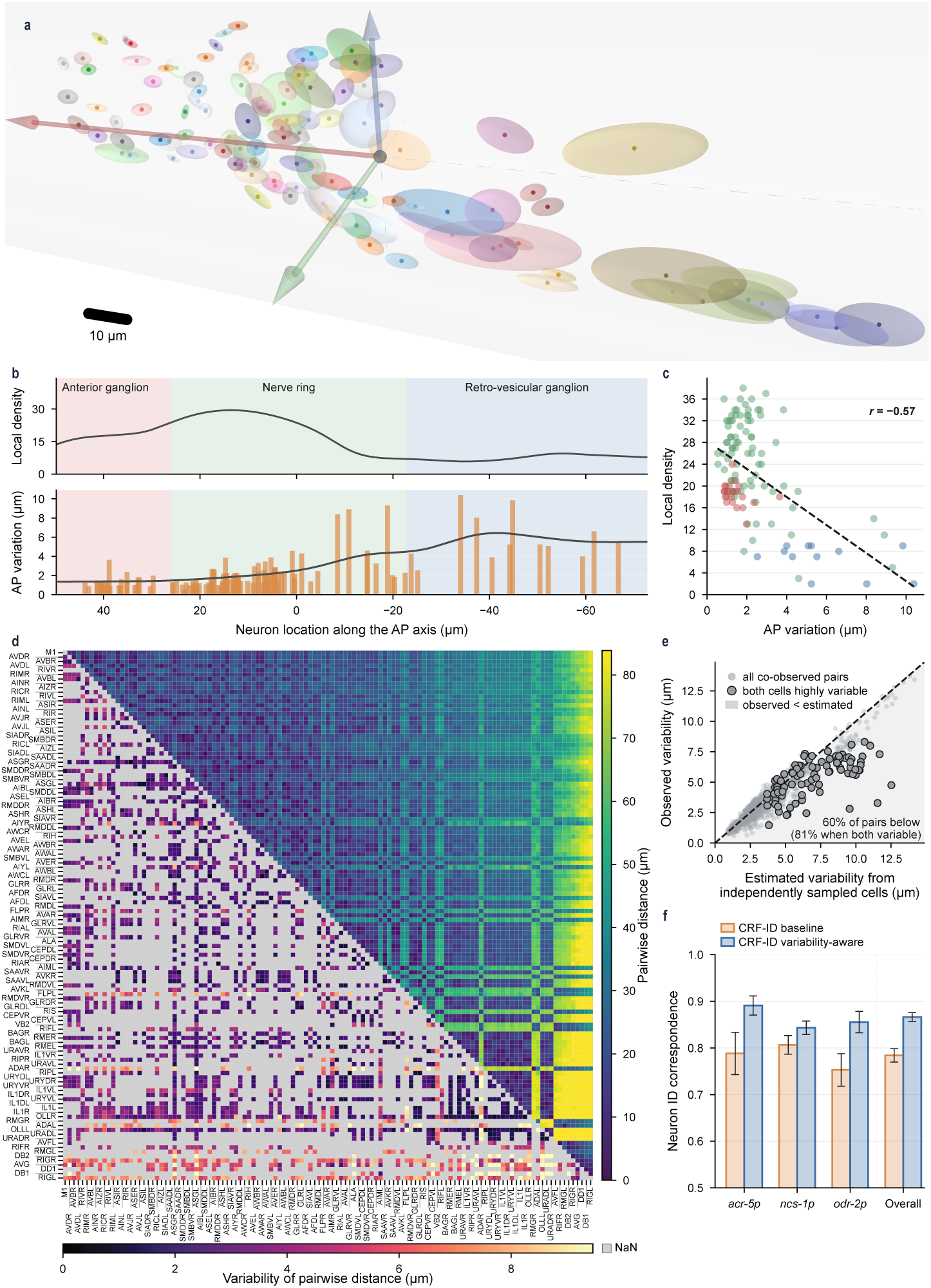
Quantified positional variability confirms the robustness of relational features and improves automated neuron identification. **a.** Positional variability of covered head neurons, shown as ellipsoids in the atlas coordinate system (arrows: anterior-posterior, red; left-right, green; dorsal-ventral, blue). Ellipsoid axes correspond to one standard deviation, N = 106 worms. **b.** Local neuron density (top) and positional variability (bottom, one bar per neuron) along the anterior-posterior (AP) direction as a function of position along the AP axis, with the anterior ganglion, nerve ring, and retrovesicular ganglion shaded. **c.** Local density versus AP variability for each neuron, colored by region as in b. Dashed line, linear fit; Pearson *r* = −0.57; two-sided *P* = 3.0 × 10*^−^*^11^; N = 117 neurons (more than 7 occurrences). **d.** Binary feature matrix for all covered neuron pairs. Upper triangle, mean pairwise distance (*µ*m); lower triangle, standard deviation of pairwise distance across worms (*µ*m); gray, pairs never co-observed. **e.** Observed variability of pairwise distance (s.d. of distance across worms) versus the variability expected if the two neurons moved independently (s.d of the distance between 1000 independent draws from each neuron’s fitted 3D Gaussian). Gray, all co-observed pairs; outlined, the 88 pairs in which both neurons are highly variable (above 70 percentile, 4.5 *µ*m); shading, observed below expected; dashed line, equality. **f.** Leave-one-out identification accuracy of the CRF-ID baseline (CRF-ID 2.0 [34]; all edges weighted equally) and of variability-aware CRF-ID (edges weighted by their reliability across atlas animals and neurons by their positional variability), for three strains and for all five strains pooled (Overall). Bars, mean; error bars, SEM.; N = 20 (*acr-5p*), 25 (*ncs-1p*), 20 (*odr-2p*) and 106 (Overall) animals. Two-sided Wilcoxon signed-rank test, *P* = 0.0031, 0.031, 0.011 and 1.2 × 10*^−^*^9^, respectively.

Relational features tell a different story. The pairwise distance matrix and its variability across animals (**Fig. 4d**, upper and lower triangles; **Supplementary Fig. 20 and 21**) show that many neuron pairs vary far less in their separation than either member does in absolute position. To ask whether this reflects biology instead of a statistical artifact of averaging, we compared the observed variability of each pair’s distance with the variability expected if the two neurons moved independently, estimated by resampling each neuron’s positions without regarding the others’ (**Fig. 4e**). Observed variability falls below the independent expectation for most pairs, and the gap is largest for pairs in which both neurons are individually highly variable. Neurons that move substantially tend to move together with their neighbors, making their relative positions more stable and reliable for identification than their absolute positions. This is the empirical basis for the relational representation described earlier, and it is consistent with the robustness of pairwise features observed in earlier identification work [22, 34].

This observation suggested a direct use for the variability estimates. In the previous CRF-ID related work [22, 34], all pairwise constraints are considered as equally important. We modified its cost function so that each constraint in the graph is weighed by the atlas-derived variance, so that stable relationships contribute more to the inference and unstable ones less. On the same atlas and the same held-out worms, the variability-aware version improves overall leave-one-out correspondence from 0.78 to 0.87, with gains in each strain tested (**Fig. 4f and Supplementary Fig. 22**). The variability that our brain-wide atlas measures is therefore not only a description of anatomy but also a usable signal for the identification task, in which we show various applications in the following sections.

### A hermaphrodite atlas supports accurate neuron identification in male worms

To further test the robustness of our atlas and overall approach, we applied our atlas to the identification of neurons in male *C. elegans*. The direct translatability of hermaphrodite atlases to male identification is complicated by several key factors. First, males differ in body dimensions. Second, males have additional sex-specific neurons. Lastly, even the position of sex-shared neurons may differ. To test robustness of our atlas to these factors, we imaged and manually annotated males of the *glr-1p*::mCherry multi-cell strain (**Fig. 5a**). In addition to identifying neurons in male worms with our atlas constructed from hermaphrodite data, we compared our results to two other conditions: performance of the hermaphrodite atlas on hermaphrodite data (hermaphrodite LOO) and performance of a male-specific atlas on male data (male LOO). For the latter two, we used the same leave-one-out scheme as previously, where predicted worms were not included in the atlas used to make their own prediction. The hermaphrodite atlas identified male neurons as accurately as the male-specific atlas did, and the difference between the two was not significant (**Fig. 5b**). The neurons the two sexes share showed relationships close enough to their hermaphrodite counterparts, that the hermaphrodite atlas was sufficient to provide accurate identities of neurons in male for this strain. We then asked how many hermaphrodite worms are needed to reach this accuracy. Building the hermaphrodite atlas from progressively more worms and applying it to the male data, the accuracy rose steeply up until about seven animals and plateaued, staying within a few points of the male leave-one-out ceiling (**Fig. 5c**). Therefore, we were able to conclude that a hermaphrodite atlas of a modest size (n = 7-10) is sufficient for obtaining high accuracy in neuron prediction for males.

**Fig. 5:**
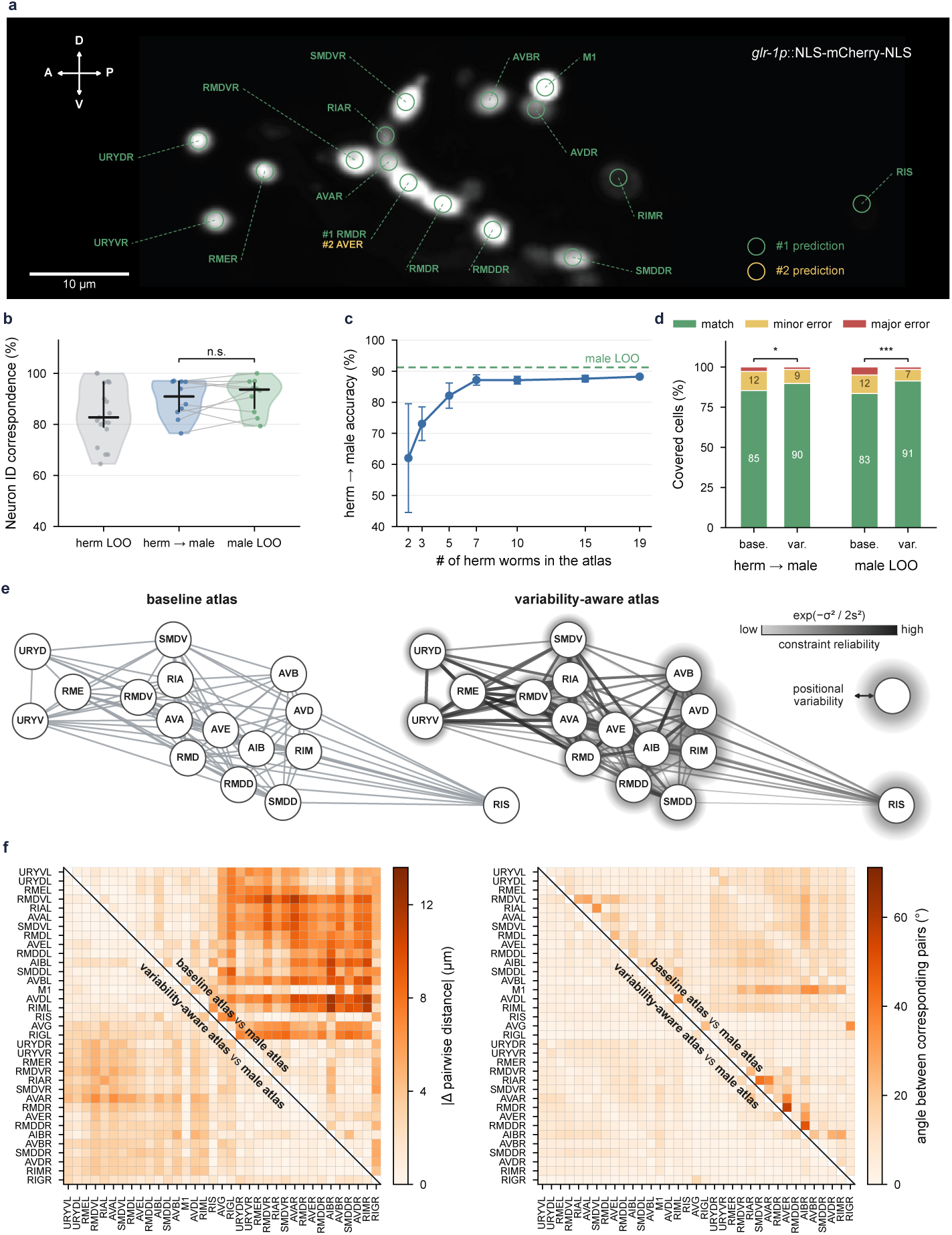
A hermaphrodite atlas enables accurate neuron identification in male worms. **a**. Representative image of male head (right half, maximum intensity projection) expressing *glr-1p*::NLS-mCherry-NLS with identities assigned by CRF-ID using the hermaphrodite *glr-1p* atlas. Green, rank-1 prediction; orange, rank-2 prediction. More examples can be found in **Supplementary Fig. 23**. **b**. Identification accuracy for three conditions: hermaphrodite atlas applied to hermaphrodite worms by leave-one-out test (herm LOO), hermaphrodite atlas applied to male worms, and male-specific atlas applied to male worms by leave-one-out test (male LOO). Violins, distribution; bars, median and interquartile range; gray lines connect the same male worm; n.s., two-sided Wilcoxon signed-rank test, *P* = 0.52. **c**. Accuracy of hermaphrodite-to-male identification as a function of the number of hermaphrodite worms used to build the atlas. Points, mean over 10 randomly sampled worms to build an atlas at each atlas size ; error bars, s.d. across atlases at the chosen size; rightmost point, the full atlas with all 20 worms; dashed line, male leave-one-out accuracy. **d**. Fraction of covered cells scored as match, minor error, or major error for hermaphrodite-to-male and male leave-one-out inference, with equal edge weights (base.) and variability-aware weights (var.). Minor error, overlapping 2-s.d. position ellipsoids of the ground truth labels. Two-sided Wilcoxon signed-rank test; \**P <* 0.05, \*\*\**P <* 0.001; N = 10 male worms. **e**. Relational graph of the *glr-1p* neurons in the baseline atlas (left; all edges weighted equally) and in the variability-aware atlas (right), built from 20 hermaphrodites (lateral view, anterior left, dorsal up). Edges join pairs annotated together in at least 5 worms; In the variability-aware atlas, edge shade and width indicate constraint reliability, exp(−*σ*^2^*/*2*s*^2^), where *σ* is the s.d. across worms of the 3D distance between the two neurons and *s* is the median *σ* over all atlas pairs co-observed in at least 5 worms (2.71 *µ*m); the halo around each node extends one s.d. of its position (mean of the AP and DV s.d.) beyond the node circle. The variability atlas for other strains are shown in **Supplementary Fig. 24**. **f**. Differences between the male and hermaphrodite atlases for the *glr-1p* neurons annotated in at least 5 worms of each sex (ordered anterior to posterior): pairwise distance between mean positions (left; |Δ|, *µ*m) and angle between corresponding pair vectors after rigid alignment of the two atlases (right; degrees). Upper triangle, difference as seen by the baseline atlas (unweighted); lower triangle, difference as seen by the variability-aware atlas, weighted by the reliability of the pair’s distance (as in **e**) and the reliability of its direction (angular s.d. of the pair vector across worms, scaled by its median). Color scale, 0 to twice the perturbation that lowers identification accuracy by 5 % in Fig. 3 (6.8 *µ*m; 35.6*^◦^*), marked at the color-bar midpoint.

The resulting prediction errors were classified based on matches: minor errors in which the predicted neuron is an anatomical neighbor of the correct one, and major errors (**Fig. 5d**). Minor errors are tolerable for many downstream uses because they place the cell in the correct neighborhood. Under both conditions, the variability-aware weighting introduced in the previous section reduced errors further, raising the match rate from 85% to 90% for hermaphrodite-to-male inference and from 83% to 91% for the male-specific atlas. The relational graphs of the shared neurons, with edges colored by pairwise distance and weighted by its variability, are closely similar in the two atlases (**Fig. 5e**). This is also shown in **Fig. 5f**, where the difference in pairwise distances and angles between neurons are similar for both atlases. With additional experimental optimization to reduce positional variability from compression when imaging with agar pads, it may be possible to further close the remaining gap observed in our atlases.

### The brain-wide atlas assigns identities in whole-brain data across imaging modalities

We evaluated the brain-wide atlas in three whole-brain settings that differ in imaging modality, the ground truth available, and in what a user would do with the identities. The first is the OpenWorm coordinate set [33], used above as a comparison atlas and here as a target: it derives from electron-microscopy reconstruction, a modality that shares no optical characteristics with the fluorescence data the atlas was built from, and every neuron in it carries an identity, so the ground truth is unambiguous. We applied CRF-ID to this point set using the brain-wide atlas (**Supplementary Fig. 25**) and scored each prediction as a match, a minor error in which the predicted label is an anatomical neighbor of the true one, or a major error (**Fig. 6a**; cells outside atlas coverage were excluded from scoring). Top-1 accuracy was 54%, rising to 86% when the three highest-ranked candidates were considered and to 92% at rank five. The gap between rank one and rank three indicates that the atlas often includes the correct identity among its top candidates even when its highest-ranked assignment is incorrect, making the candidate list particularly useful for users who manually review predictions.

**Fig. 6:**
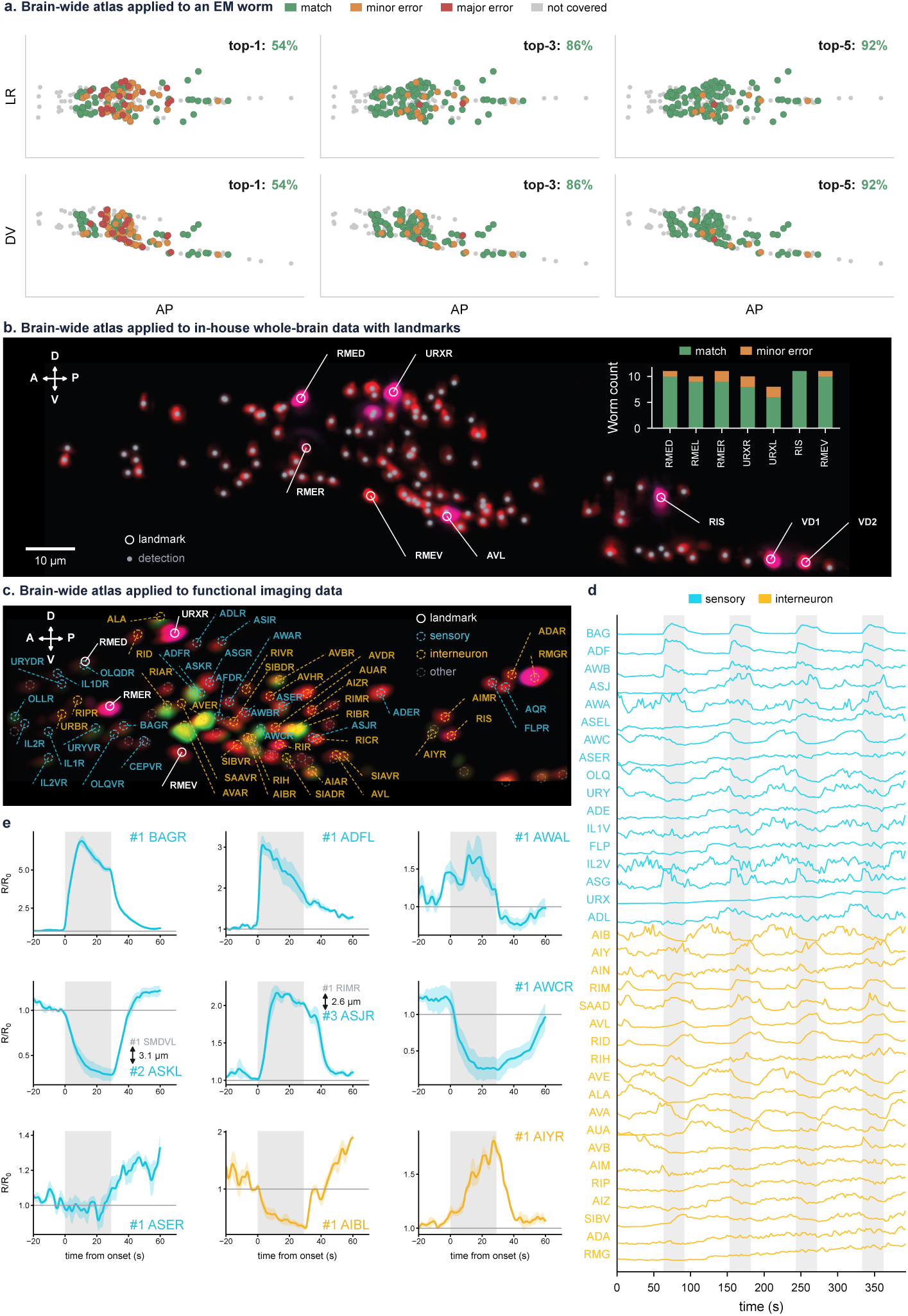
The brain-wide atlas assigns identities in whole-brain data across imaging modalities. **a**. Application of brain-wide atlas to prediction of neuron identities from the OpenWorm neuron coordinate set [33], which derives from electron-microscopy reconstruction, shown in left-right (top) and dorsal-ventral (bottom) views. Green, match; orange, minor error; red, major error; gray, not covered by the atlas. Top-1, top-3, and top-5 accuracy are given for each column. **b**. A representative image (right half, maximum intensity projection) of pan-neuronal GT408 strain (red, pan-neuronal marker; magenta, CyOFP landmark cells indicated by white circles; gray points, detected nuclei). Inset: number of worms in which each landmark cell was assigned its correct identity (green), a neighboring identity (orange), or a non-neighboring identity (red), across 10 animals. Scale bar, 10 *µ*m. **c**. Whole-brain functional imaging GT408 worm (right half, maximum intensity projection) with identities assigned from the red channel by the brain-wide atlas. White circles, landmark cells; cyan, cells assigned sensory identities; orange, interneuron identities; gray, other. Scale bar, 10 *µ*m. **d**. GCaMP traces of a subset of identified sensory and interneuron cells in **c** over the full recording with tracking done by [36]; shaded intervals, OP50 stimulus. **e**. Trial-averaged responses (*R/R*_0_; line, mean; shading, SEM across trials aligned to stimulus onset (gray, OP50 pulse) for representative cells; labels give the top-ranked atlas candidate (#1). When the #1 prediction’s activity deviates from what was reported [37], the top-ranked candidate is shown in gray and an arrow points to the matching candidate in terms of activity with its rank (#*k*; cyan, sensory neuron; yellow, interneuron), next to the distance from the #1 to #*k* candidate’s mean atlas position.

The second use-case is the one the atlas was most directly built for – pan-neuronal fluorescence in a single channel, where manual annotation is exceptionally difficult and is the problem that multicolor strategies were designed to solve. To characterize the performance of our atlas and pipeline, we used a strain carrying pan-neuronal mNeptune, pan-neuronal GCaMP6s, and a CyOFP landmark set [22, 38]. Nuclei were detected in the mNeptune channel and identities were assigned with the brain-wide atlas (**Fig. 6b**). The landmark cells provide an internal benchmark because their identities are independently established by the landmark channel, allowing predictions to be evaluated in every animal without annotating the full volume. Across 10 animals, the atlas assigned correct identities to the covered landmark cells in most worms, with the residual errors concentrated in confusions between members of the same anatomical class (**Fig. 6b**, inset). Landmark cells that lie outside the coverage of the head atlas were, as expected, not recovered reliably, since the atlas holds no positional information for them. Landmark-based scoring of this kind requires no manual annotation and can be applied by any user to their own recordings as a per-animal quality check.

The third use case examines neuronal identification in whole-brain functional imaging, where matching neuronal identities across animals is essential for comparing neural activity. We recorded calcium activity in an animal expressing pan-neuronal GCaMP with the same landmark set, assigned identities from the red channel with the brain-wide atlas, and grouped the identified cells into sensory and interneuron classes according to their predicted labels (**Fig. 6c**). Neural activities across the identified population are shown over the full recording, with four presentations of OP50 supernatant marked (**Fig. 6d**). Because responses to a defined stimulus are constrained by known physiology, the traces provide a sanity check on the assignments that are independent of any manual annotation. Trial-averaged responses aligned to stimulus onset are shown for representative cells together with the three highest-ranked candidate identities from the atlas (**Fig. 6e**). Cells assigned to sensory classes known to respond to the stimulus showed increases in fluorescence time-locked to onset. Cells assigned to AWC and to its downstream partner AIB show decreases during the stimulus and recovery at its offset, consistent with the established inhibition of AWC by attractive odorants and the excitatory connection from AWC to AIB [39]. The agreement between predicted identity and expected response across the examined cells indicates that the assignments are not arbitrary, and it shows that identities supplied by the atlas are sufficient to interpret whole-brain activity at the level of individual neurons.

Response profiles can also adjudicate among the candidates the atlas returns. For instance, in one cell, the highest-ranked assignment was RIMR, but the observed response matched the physiology expected of the third-ranked candidate, ASJR (**Fig. 6e**). This example illustrates how functional signatures, which provide information independent of neuronal position, can distinguish candidate classes when their responses to a stimulus are known. This is consistent with how neuronal identities are assigned manually, where researchers often combine anatomical and physiological evidence. Together, these findings suggest a workflow in which the atlas narrows identity to a small set of candidates, with physiological criteria resolving the remaining ambiguity without requiring the top-ranked atlas assignment to be correct.

## Discussion

We presented a pipeline for building data-driven, anatomically faithful, continuously updatable brain-wide atlases of the *C. elegans* nervous system at single-neuron resolution. The atlas is assembled from partially overlapping multi-cell strains, integrated through anatomy-anchored alignment and probabilistic consensus annotation, and represented as a graph in which neurons are nodes and pairwise positional relationships are edges. Three features distinguish our approach from previous atlas-building methods: the use of relative rather than absolute neuronal positions, anatomically defined alignment axes independent of neuronal labels, and a graphical representation that allows the atlas to be extended without rebuilding it. To the best of our knowledge, this is the first single-neuron-resolution anatomical atlas of a nervous system designed to be continually refined and expanded as new data become available.

Our approach supports both direct application of existing atlases and construction of atlases tailored to specific experimental conditions. Users can apply the atlas to their own data without acquiring or crossing our strains, while laboratories studying particular strains or conditions can construct matched atlases that substantially outperform a generic one. Atlas construction requires only a single fluorophore, a standard confocal microscope, and a microfluidic device fabricated by soft lithography, making it more accessible than multicolor approaches. Moreover, the atlas can be incrementally extended rather than rebuilt, allowing laboratories to contribute data collected under their own conditions to a shared atlas while maintaining context-specific atlases for their experiments.

Beyond its use as a reference for identification, the atlas is itself a measurement of positional variability and structure in these variations. Across many individuals, the positional variability of head neurons is non-uniform with neurons in densely packed regions varying less along the anterior-posterior axis than those in sparsely populated regions, consistent with mechanical constraint imposed by their neighbors. Relational features further reveal that neighboring neurons tend to move together, making pairwise distances more stable than absolute positions. Consequently, weighting these spatial relationships by their measured variability improves automated neuron identification (**Fig. 4e and f**). The same measurements help explain where human annotation is hard, although mosaicism in marker expression predicts inter-annotator disagreement more strongly than positional variability alone (**Supplementary Fig. 19**). While these observations are descriptive and not mechanistic, they provide a measured baseline for work on the developmental robustness of neuronal positioning.

The atlas in its current form has limitations that bound where it can be applied. Coverage is approximately 66% of head neuron classes; the gap reflects the promoters available in our current panel. However, the design of our atlas also makes this easily addressable, due to the ease of incorporating new data from new strains after further promoter screening. Construction depends on the quality of human annotation, and although agreed criteria and BFP-aided verification reduce the error, residual mistakes propagate into the atlas in ways that are hard to audit. The alignment requires both pharyngeal bulbs to be detected, which fails in animals with unusual posture or with imaging artifacts in that region; such animals are currently excluded by hand. Generalization has been demonstrated in males, where the hermaphrodite atlas performed comparably to a male-specific atlas, and in whole-brain pan-neuronal recordings. However, its performance under conditions that differ more substantially from those used for atlas construction, including imaging setups, remain untested. Applications that diverge substantially in worm age, posture, or with tissue deformation may need further fine-tuning through a condition-matched extension instead of direct application of the existing atlas.

We envision several uses of the pipeline. The atlas can be extended to different genetic backgrounds, developmental stages, sexes, and imaging modalities. Once several condition-specific atlases exist, they can be compared directly, asking which positional features are conserved across genetic backgrounds and which are not; the framework that supports inference also supports that comparison. Integration with functional imaging using GCaMP is particularly attractive because the green channel remains free in our minimally loaded strains, so the activity and atlas-derived identity can be recorded in the same animal. Beyond *C. elegans*, the combination of relational representation, anatomy-anchored alignment, and incremental extension may suit other systems in which individual variability and sparse labeling together defeat coordinate-based approaches. The broader claim we would make, beyond the *C. elegans* atlas itself, is that the trade-off between fidelity, efficiency, and accessibility is a consequence of design choices.

## Methods

### Strains and maintenance

*C. elegans* strains were maintained at room temperature (∼ 22 *^◦^*C) on 6 cm NGM (nematode growth medium, 1.6 % agar, 2.5 g/L Bacto Peptone, 3 g/L NaCl, 25 mM KPO_4_ at pH 6.0, 1 mM CaCl_2_, 1 mM MgSO_4_, 5 mg/L cholesterol) plates containing *E. coli* OP50 as food source by following standard protocols [40]. Strains were passaged by transferring a small piece of agar containing worms or by transferring around 20 L4-stage hermaphrodite larvae from a well-grown plate to a fresh NGM plate containing a lawn of *E. coli* OP50. Strains were maintained under the standard condition for at least two generations before being used for testing behavioral or other traits.

### Generation of transgenes and transgenic animals

The transgenes that express a nuclear localized mCherry reporter in subsets of neurons were previously described and published [29]. The transgenes that express a nuclear localized blue fluorescent protein (NLS-BFP) in selective neuronal cell types to verify cell identification were generated using LR recombination (Invitrogen #11791020) as previously described [29]. The transgenic worm strains stably expressing the nuclear localized mCherry reporter transgenes were generated by injecting transgenes, and co-injection DNA markers when specified, at concentrations ranging from 10 to 200 ng/µL using the standard method [41].

### Synchronization and staging

Animals were synchronized by timed hatch-off. Four to five L4 animals were placed on a fresh plate and allowed to lay eggs for approximately four days, until the plate was densely covered with eggs while the bacterial lawn remained visible. Adults and larvae were rinsed off with S-basal, and the remaining eggs were allowed to hatch for one hour. Hatched L1 larvae were collected in S-basal, pelleted by brief centrifugation, and transferred to a 100 mm seeded plate. Animals were grown at 20 *^◦^*C and imaged 62 to 63 h after hatch-off, as day-1 adults; the one-hour spread reflects consecutive imaging sessions from the same synchronized population.

### Microfluidic device fabrication

The array-style imaging device was adapted from a previous design from our laboratory [42]. The main modification was the removal of channel restrictions near the side-channel inlets, which prior loading experiments had identified as a source of inconsistent orientation: the restriction forced adult animals to rotate as they entered the imaging channels. With the restriction removed, the narrowest point of each channel lies within the imaging region itself. Each device carries seven rows of approximately 20 side channels. Channels are 65 *µ*m wide and 60 *µ*m high, with a single uniform height across the device, and each terminates in an opening 8 to 10 *µ*m wide that allows fluid to pass while retaining the animal. A variant with 60 *µ*m-wide channels is used for animals imaged before the 62 h timed hatch-off.

Device designs were drawn in CAD and printed as photolithography masks by a commercial vendor. Masters were fabricated from the masks by standard SU-8 photolithography in a cleanroom. After silanization of the master, polydimethylsiloxane (PDMS; Sylgard 184) was mixed at a 10:1 ratio of base to curing agent, degassed under vacuum, poured onto the master, and cured overnight at 65 *^◦^*C. Cured PDMS was peeled from the master, punched for inlet and outlet ports, and bonded to glass coverslips after oxygen-plasma treatment. Tubings were rinsed with deionized water after each session and reused.

### Worm loading and image acquisition

Tubing was assembled from 1/32-inch inner-diameter silicone tubing and 19-gauge blunt syringe adaptors, primed with S-basal filtered through a 0.22 *µ*m membrane, and connected to the device inlet and outlet. The device was mounted on a Petri dish, degassed by closing the outlet and applying gentle syringe pressure until air bubbles cleared, and inspected for leaks. Synchronized adults were transferred from the hatchoff plate into a microcentrifuge tube, allowed to settle, drawn into the inlet tubing, and pushed into the device by hand with a 3 mL syringe. Under these conditions, at least 100 animals entered the imaging channels within approximately three minutes. After loading, 0.5 mL of 50 mM sodium azide in S-basal was introduced in the device to immobilize the animals. Finally, after both clamps were closed, the device was transferred to the microscope for imaging.

Volumetric images were acquired on a Nikon CSU-W1 Yokogawa spinning disk field scanning confocal microscope equipped with a Hamamatsu ORCA-Fusion Gen-III sCMOS camera. Multichannel Z-stack fluorescence images were taken using a 40× oil-immersion objective (NA 1.4). mCherry was excited and collected with the manufacturer’s default mCherry settings (laser line 561 nm; emission filter 620 W60) at 100 % laser power for *acr-5p* and 60 % laser power for the other strains. Brightfield images were also acquired at each plane for landmark detection. Stacks were acquired with a lateral pixel size of approximately 0.15 *µ*m and a z-step of 0.3 *µ*m, spanning 160 to 180 optical sections at 100 ms exposure per section. Only animals lying on their side, with the head fully within an imaging channel and no visible compression, were selected for acquisition; roughly half of the loaded animals met this criterion. A summary of the datasets used to build the atlas is provided in **Supplementary Fig. 13**.

### Confocal microscopy for cell identification in males

Worms were maintained at room temperature (∼ 22 *^◦^*C) using the standard procedure described above. To prepare male worms, 10 transgenic hermaphrodites were crossed with 15 – 20 wild-type males. 30 – 50 L4-stage transgenic male progenies from the cross were transferred to a fresh NGM plate and kept at room temperature overnight. Confocal microscopy for acquiring male data was conducted using a Nikon Eclipse Ti-E inverted confocal microscope equipped with a Yokogawa CSU-X1 spinning disk scanner unit and an Andor iXon Ultra EMCCD camera. Multichannel Z-stack fluorescence images were taken using a 40× oil-immersion objective (NA 1.3) and a Piezo Z stage with a step size of 0.3 µm. A 561 nm laser and a single band fluorescence emission filter centered at 617 nm (*>* 90 % transmission over 50 nm) for taking images red fluorescence channel. For imaging, around 15 day-1 adult males were picked into a drop of 2 *µ*L of 50 mM sodium azide solution on an agar pad freshly prepared with 2 % agar (w/v, SeaKem LE Agarose, Lonza #50004). To control the thickness of the pad, the melted agar was placed on a glass slide positioned between two supporting glass slides, each with two layers of laboratory tape on top, and pressed with a glass slide placed on top. A cover glass was gently placed on the pad when worms were immobilized and ready for imaging. Images were taken using NIS Elements AR (version 4.40, Nikon) and analyzed for co-localization using ImageJ. Nuclei in the male data were detected manually and annotated independently by two experts; the consensus of these annotations was used as ground truth for the male analysis in **Fig. 5**. Male nuclei were then identified against the brain-wide atlas built from hermaphrodite data, and predictions were scored against this consensus.

### Aversive training with pathogenic bacteria

Worms were maintained at room temperature (∼ 22 *^◦^*C) using the standard procedure described above. To prepare worms for aversive training, around 60 L4-stage hermaphrodite larvae were transferred to a fresh 6 cm NGM plate containing an OP50 lawn one day before behavior experiment and kept at room temperature. To prepare NGM (nematode growth medium, 2.5 g/L Bacto Peptone, 3 g/L NaCl, 25 mM KPO_4_ at pH 6.0, 1 mM CaCl_2_, 1 mM MgSO_4_, 5 mg/L cholesterol) culture of *E. coli* OP50 or *P. aeruginosa* PA14, individual bacteria colonies were picked in 30 mL NGM medium and grown at 27 *^◦^*C with shaking at 114 rpm in an incubator shaker (BT Lab Systems) for around 12 hours or until appropriate density. To prepare aversive training or control plates, 6 – 10 mL respective culture was transferred to an empty 10 cm NGM plate to cover the entire surface of the plate by gently rotating the plate, and the excess amount of culture was removed with a pipette and used to prepare another training or control plate if needed. The plates were incubated at 27 *^◦^*C for approximately 36–48 hours. Before being used for training, the plates were placed on bench at room temperature with the lid half-open for about 30 minutes. To train worms, 15 – 20 day-1 adult hermaphrodites were transferred to each PA14-training or OP50-control plate and trained at room temperature for 4 – 5 hours before learning assay.

### Microdroplet assay for testing olfactory response and learning

Microdroplet assay was performed as previously described [43] with minor modifications to analyze odorant response and aversive learning behavior. Twenty mL of NGM culture of OP50 or PA14 prepared as described above was transferred into a glass flask with a side arm to use as the odorant source. Twelve 1.5 *µ*L droplets of NGM buffer (3 g/L NaCl, 25 mM KPO_4_ at pH 6.0, 1 mM CaCl_2_, 1 mM MgSO_4_) were placed on a sapphire window. Individual day-1 adult hermaphrodite were first briefly rinsed in NGM buffer and then transferred to the droplets with one worm in each. The sapphire window was gently placed into an enclosed chamber connected with an air stream odorized by flowing above the bacteria cultures contained in the flasks and illuminated with LEDs. The air stream flowing into the chamber alternated between the one odorized with OP50 culture and the one odorized with PA14 culture every 30 seconds for 12 cycles. The assay was recorded with a Watex camera at 10 frames per second and the locomotion of the worms was analyzed using a Matlab code [44] to identify large body bends and turns. A choice index is defined as [[(turning rate to OP50) – (turning rate to PA14)] / Total turning rate] and a learning index is defined as [(choice index of naive worms) – (choice index of trained worms)]. Training with PA14 generates a positive learning index. The same procedure was followed to measure response to OP50 when the air stream alternated between one odorized with OP50 and the other with NGM buffer. A chemotaxis index is defined as [[(turning rate to buffer) – (turning rate to OP50)] / Total turning rate]. Three worms were used in each assay for each genotype each condition, 2 – 4 replicates were performed each day which were repeated on 3 – 4 different days.

### Locomotion analysis

*C. elegans* strains were maintained at room temperature (∼ 22 *^◦^*C) using the standard procedure described above. To prepare worms for locomotion analysis, L4-stage hermaphrodite larvae were transferred to a fresh 6 cm NGM plate containing an OP50 lawn one day before the experiment and kept at room temperature. To prepare plates for recording, a small amount of NGM culture of OP50 was spread on a 6 cm NGM plate, incubated at 27 *^◦^*C for 36 hours, cooled and dried on bench for 1 hour before use.

To record worms’ locomotion on food, approximately 10 day-1 adult hermaphrodites were transferred to a plate prepared for recording and kept at the room temperature for 30 minutes before recording started. The worms on each plate were recorded for 10 minutes with a Grasshopper3 GS3-U3-120S6M camera (Point Grey) equipped with a 60 mm f/2.8D AF Micro-Nikkor lens (Nikon) at 7 frames per second and the locomotion of the worms were analyzed using Wormlab (MBF Bioscience, version 2024.1.1). Three plates each containing approximately 10 worms were recorded and analyzed each day, which were repeated on 3 different days.

### Chemotaxis assay with IAA

Chemotaxis analysis on individual worms were performed as previously described [45]. To prepare assay plates, 8 mL NGM medium was poured into each 10 cm petri dish, dried with the lid open on bench for 1 hour, and kept on bench with the lid on at room temperature overnight until use. To prepare worms for chemotaxis assay, L4-stage hermaphrodite larvae were transferred to a fresh 6 cm NGM plate containing an OP50 lawn one day before the experiment and kept at room temperature until the assay. Isoamyl alcohol (IAA) was serially diluted to 10^-3^ with ethanol. To set up the assay, a single day-1 adult worm was first transferred onto an empty NGM plate to crawl for approximately 30 seconds to remove residual bacteria from the body. Ten *µ*L IAA (10^-3^, v/v) was pipetted onto an assay plate 1.5 cm away from the center, followed by placing the worm from the transitional plate to the center of the assay plate immediately before the recording started. The recording was terminated when the worm reached IAA; otherwise, recording continued for 5 minutes. The recordings were analyzed using Wormlab (MBF Bioscience, version 2024.1.1) and the efficiency of chemotaxis was reported with Navigation index (Radial speed / Traveling speed) as previously described [45]. Three worms were assayed for each genotype every day which were repeated for 3 different days.

### Brood size measurement

The method to measure brood size was developed based on previous studies [46, 47]. *C. elegans* strains were maintained as described above. Individual L4-stage hermaphrodite worms (P0s) were transferred to fresh 6 cm NGM plates with one worm per plate. The plates were kept at room temperature for 36 hours before the P0 worms were removed [47]. 36 -48 hours later the plates were imaged using a Grasshopper3 GS3-U3-120S6M camera (Point Grey) equipped with a 60 mm f/2.8D AF Micro-Nikkor lens (Nikon) capturing each plate entirely. The number of worms on each plate was counted using Wormlab (MBF Bioscience, version 2024.1.1) and unhatched eggs were counted as dead. 10 P0 worms were measured for each genotype each day which was repeated on 3 different days.

### Neuron detection and anatomical landmarks

From the volumetric images, the nuclei were detected in the mCherry channel with a StarDist model [48] trained on in-house multi-cell volumes, and every detection was inspected and corrected manually by a single curator before annotation. All annotators therefore worked from the same set of nuclei, and no post-hoc reconciliation of nucleus sets between annotators was needed. The same procedure was applied to the pan-neuronal red channel of GT408.

The two pharyngeal bulbs, the metacorpus and the terminal bulb, were detected in the brightfield stack by a vision-transformer model trained on 50 manually annotated brightfield images. The centroid of the terminal bulb defined the origin of the worm coordinate system, and the line through the two bulb centroids defined the anterior-posterior (AP) axis. Each bulb detection was compared with a consensus reference by structural similarity (SSIM), and detections with low similarity were queued for manual inspection first; in practice the large majority required no correction (**Supplementary Fig. 9**).

### Annotation criteria, auxiliary BFP lines, and consensus annotation

Annotation criteria were established separately for each construction strain before any atlas data were labeled. For each strain, we generated auxiliary transgenic lines co-expressing BFP from cell-selective promoters on the multi-cell mCherry background, targeting neurons that annotators found ambiguous from the mCherry channel alone [29]. Co-localization of BFP and mCherry in these lines identified the anatomical features that reliably distinguish each ambiguous neuron from its neighbors, including relative position to specific neighbors, membership in a cluster, and position along an anatomical landmark and relative to the body surface. These features, together with published expression patterns [18, 32], were written into per-strain criteria (**Supplementary Fig. 6**) that were reviewed and agreed by all annotators before labeling. The BFP lines were used only for this purpose and did not contribute to positional data for the atlas.

Each volume was then annotated by three to four independent annotators. For each nucleus, the annotator recorded a single identity and a confidence score from 0 to 3, where 3 denotes high confidence and 0 denotes that no identity could be assigned. Annotators worked without access to each other’s labels. After independent annotation, the annotators met to reconcile disagreements worm by worm using the annotation interface (**Supplementary Fig. 7**), which displays every annotator’s label for each nucleus with the annotators’ names hidden. Most disagreements resolved to a shared identity on discussion; where they did not, the consensus was recorded with a reduced confidence score or, if no identity could be defended, left unassigned. Individual annotations were retained alongside the consensus. The timeline for annotation from scratch was estimated by the experience across approximately 100 volumes and three to four annotators, not the result of a timed comparison (**Supplementary Fig. 4**).

### Two-stage anatomy-assisted alignment

Annotated centroids from each volume were expressed in the worm coordinate system defined by the terminal bulb origin and the AP axis. With these two features fixed, the only remaining degree of freedom per animal is rotation about the AP axis. Alignment across datasets proceeded in two stages.

In the first stage, alignment was performed within each strain. Animals were aligned sequentially: each new animal was rotated about the AP axis by the angle that minimized the sum of squared distances between its labeled nuclei and the corresponding nuclei of the animals already aligned, found with a gradient-based optimizer. After all animals of a strain were aligned, the positions of each labeled neuron were pooled to give a per-neuron positional distribution.

In the second stage, alignment was performed across strains, operating on the per-neuron distributions rather than on individual animals. For each pair of strains with shared neurons, a rotation about the AP axis was chosen to bring the shared neurons’ distributions into agreement in both mean and variance. The origin and AP axis remained fixed across all strains. Strains were combined greedily: starting from an arbitrary strain, the strain with the largest overlap with the current combined atlas was added next, and so on. Because the order of addition is determined by overlap rather than chosen in advance, the resulting atlas does not depend on the starting strain.

The two-stage procedure was compared with a registration-only baseline in which all animals were aligned by rigid registration on their shared labeled centroids, without the anatomical origin and axis. We randomly picked a starting worm as the reference and calculated the run-to-run variation for the two methods, and as the results shown in **Supplementary Fig. 11**, the proposed two-stage is substantially more robust, reaching the same statistical atlas regardless of which worm one starts with.

### Atlas representation and feature extraction

The aligned data were stored as a graph in which each neuron class is a node and each pair of classes is an edge. Each node carries the unary features of its neuron: the mean position across aligned animals and the standard deviation along the AP, left-right (LR), and dorsal-ventral (DV) axes, obtained by fitting a three-dimensional Gaussian to the aligned centroids. Each edge carries the binary features of its pair: the vector connecting the two neurons in each aligned animal, from which the pairwise distance (vector magnitude) and its variability across animals are derived. The direction of the connecting vector is stored as well; angles are not stored as atlas features but are computed at comparison time as the angle between corresponding vectors in two atlases or between an atlas and a target dataset. The graph is fully connected across covered neurons. For pairs never observed together in a single animal, the connecting vector is computed from the aligned mean positions of the two neurons in the combined atlas. The atlas covers approximately 66 % of head neuron classes (55 out of 83), giving 7,750 pairwise edges, of which 2,537 (33 %) were directly co-observed in at least one strain.

### Comparison atlases

Two positional references from outside this work were used as comparisons. An Open-Worm atlas was built from neuron coordinates provided by the OpenWorm project on request [33]; because these coordinates come from a single reconstruction, the atlas is static and carries no positional variability. A NeuroPAL atlas was built from the positional atlas published with NeuroPAL [11], restricted in each test to the neurons labeled by the multi-cell strain under evaluation. Both sources cover all head neurons, so no neuron labeled in a construction strain was missing from either.

### Validation with the held-out strain

Matched-strain accuracy was assessed by leave-one-out within each construction strain: each animal was held out, an atlas was built from the remaining animals of the same strain, and the held-out animal’s nuclei were identified against that atlas and against the two comparison atlases (**Fig. 3a**). Accuracy was scored as described under *Automated identification*.

For the held-out-strain test, the brain-wide atlas was built from the five construction strains with *flp-3p* excluded, and *flp-3p* animals were identified against it. A *flp-3p*-specific reference was built by leave-one-out within *flp-3p*. The four SIB neurons labeled by *flp-3p* are not present in any construction strain and were excluded from all *flp-3p* scoring.

To determine the tolerance of identification to differences between atlases, every pairwise distance, or every pairwise angle, in the *flp-3* -specific atlas was shifted by an amount drawn from a zero-mean Gaussian, leave-one-out identification was repeated, and accuracy was recorded as a function of the Gaussian’s standard deviation; the perturbation at which accuracy fell by five percentage points was taken as the tolerance (**Fig. 3f**). The observed difference between the brain-wide and *flp-3p*-specific atlases was computed per pair as the absolute difference in pairwise distance and as the angle between corresponding pair vectors; pairs were classified as co-observed if both neurons were labeled together in at least one construction strain and as inferred otherwise (**Fig. 3g–i**).

### Positional variability analysis

Positional variability was summarized per neuron as the standard deviation of aligned position along each axis and displayed as an ellipsoid with semi-axes equal to one standard deviation (**Fig. 4a**). Local neuron density along the AP axis was estimated by kernel density (Gaussian-weighted local mean) over neuron mean positions, and the correlation between AP variability and local density across neurons was assessed by Pearson correlation (**Fig. 4b and c**).

To test whether the variability of a pairwise distance is smaller than expected if the two neurons moved independently, we used every pair of neurons co-observed in at least 15 animals of one strain (1,006 pairs). Observed variability was the standard deviation of the pair’s distance across those animals. For the independent expectation, each neuron’s positions in the same animals were fitted with a 3D Gaussian (mean and covariance), 1,000 positions were drawn independently for each neuron, the pairwise distance was computed for each draw, and its standard deviation was taken as the expected variability. Both neurons of a pair were considered highly variable when the positional variability of each (root-mean-square 3D deviation from its strain mean) exceeded 4.5 *µ*m, the 70 percentile over pair members (88 pairs; **Fig. 4e**).

Annotator confusion was computed for each neuron of each strain with at least four detections (162 neurons) as the fraction of annotator pairs that assigned different labels to the same nucleus, averaged over the neuron’s nuclei, 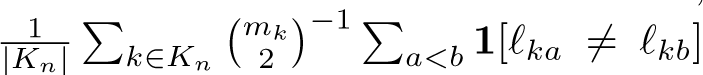, where *m_k_* ≥ 2 annotators labelled nucleus *k*. Cell-count noise was the absolute difference between the number of other nuclei detected within 8 *µ*m of each nucleus in its animal and the number expected, the count of the strain’s other atlas neurons whose mean positions lie within 8 *µ*m of the neuron’s mean position, averaged over the neuron’s nuclei. Spearman correlations between confusion and each candidate source were computed across neurons, with 95 % confidence intervals from 2,000 bootstrap resamples of neuron classes **Supplementary Fig. 19**.

### Automated identification

Neuron identities were assigned with CRF-ID, a conditional random field framework developed previously in our laboratory [22, 34]. The target dataset is represented as a graph over detected nuclei; the cost of a candidate labeling combines unary terms, comparing each nucleus’s position with the atlas distribution of its candidate identity, and with binary terms, comparing each pair of nuclei with the atlas relationship between their candidate identities. Inference finds the labeling that minimizes total cost by loopy belief propagation [49], and returns a ranked list of candidate identities for each nucleus.

The variability-aware variant uses the atlas variability in three ways. (i) Each identity pair (*i, j*) gets a reliability 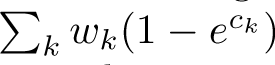, where *σ_ij_* is the standard deviation of the distance between *i* and *j* across the atlas animals in which both were annotated and *s* is the median of the positive *σ_ij_*; pairs co-observed in fewer than two animals get *T_ij_* = 0. The binary log-potential of nuclei (*a, b*) for identities (*i, j*), 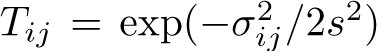, is multiplied by *f* + (1 − *f*) *T_ij_* with *f* = 0.5, so stable relationships contribute more than variable ones. (ii) In loopy belief propagation, the messages of each binary factor are raised to the power ∑ *_ij_P_ab_*(*i, j*) *T_ij_*, where *P_ab_* is the factor’s normalized potential. (iii) A unary term scores identity *i* for a nucleus at aligned position **x** by 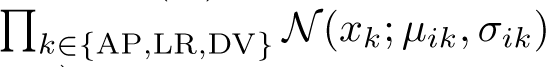, normalized over identities (the baseline unary is uniform). In addition, each nucleus is connected to its 10 nearest nuclei rather than to all others, the distance term is the squared relative difference 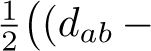 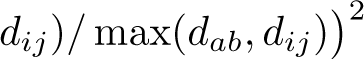, and belief propagation runs for up to 50 rather than 10 iterations; the feature weights (1, 1, 1, 0.2, 5) are unchanged.

Accuracy was scored per animal as neuron ID correspondence, the fraction of manually annotated nuclei whose top-ranked predicted identity matched the consensus label; top-*k* accuracy counts a nucleus as correct if the consensus label appears among the *k* highest-ranked candidates. For the error-severity analyses (**Fig. 5d and 6a; Supplementary Fig. 15**) a prediction was scored as a minor error if the predicted identity is a nearest neighbor of the correct one in the atlas and lies within two standard deviations of the correct neuron’s positional variability, and as a major error otherwise.

### Whole-brain applications

#### OpenWorm point set

As a target dataset unrelated in modality to the fluorescence data the atlas was built from, we used the OpenWorm neuron coordinates described above [33], which derive from electron-microscopy reconstruction and carry an identity for every neuron. The coordinates were expressed in the worm coordinate system by first using an isotropic scaling factor to globally scale and then rigidly aligned to our brain-wide atlas, identification was performed with the brain-wide atlas, and predictions were scored against the OpenWorm identities (**Fig. 6a**).

#### GT408 landmark benchmark

GT408 worms (*rgef-1p* nuclear GCaMP6s and mNeptune and *unc-47p*+*gcy-32p* CyOFP) were age synchronized and imaged using procedures described in previous section, and the images were acquired with a ZEISS Lattice SIM 3 using a 488 nm laser with a dichroic LP 560 filter to image CyOFP, and a 561 nm laser with a dichroic BP 490-560/LP 640 filter to image mNeptune. Nuclei were detected in the mNeptune channel and identified with the brain-wide atlas. Because the landmark cells are well separated, their identities were assigned by visual inspection of the CyOFP channel in the annotation interface and used to score the prediction at each landmark nucleus in each animal (**Fig. 6b**).

#### Functional imaging

Functional whole-brain imaging was performed by adapting procedures described previously [38]. Briefly, age-synchronized GT408 (day 1 adults, 70-80 hours post hatching) were imaged at 3.3 volumes per second using an open-top light sheet microscope. Each volume contained 60 slices spaced approximately 1.25 *µ*m apart, with the GCaMP6s, CyOFP, and mNeptune (green, red, and far red) channels acquired simultaneously. The odor stimulus applied was OP50 supernatant, which was prepared by centrifuging a fresh overnight culture of OP50 in LB broth at 3,000 RCF for 10 minutes. The stimulus sequence was 60 s of NGM buffer, followed by 30 s of OP50, repeated for four cycles. Stimulus delivery and timing were validated by adding 2 *µ*M fluorescein to the stimulus stream. Nuclei were tracked over time with ZephIR, and identities were assigned from the mNeptune channel with the brain-wide atlas. Calcium traces were computed per nucleus as the ratio of GCaMP6s to mNeptune fluorescence, and *R/R*_0_ was obtained by normalizing to the mean of the dimmest 10 % of time points [38]. Trial-averaged responses were computed by aligning the four presentations to stimulus onset. Cells were grouped as sensory or interneuron according to the class of their top-ranked predicted identity.

## Software and hardware

Nuclei were detected with StarDist (0.9.2) [48]; the annotation interface was built on napari (0.5.5); identification used our Python implementation of CRF-ID [22, 34] with the variability-aware extensions described above. Analyses used Python 3.12.7 with NumPy 1.26.4, SciPy 1.13.1, pandas 2.2.2, Numba 0.60.0, scikit-image 0.24.0, scikit-learn 1.5.1, statsmodels 0.14.2, tifffile 2023.4.12 and Matplotlib 3.9.2. The microfluidic device was designed in AutoCAD. CRF-ID ran on the CPU of a laptop (8 cores, 16 GB RAM, Windows 10) and took about 1 s per animal (median 1.1 s for the variability-aware and 1.3 s for the baseline CRF-ID).

## Data and code availability

The YAtlas App and the demo for CRF-ID are available at https://github.com/lu-lab/YAtlas. The raw image volumes, the annotated point clouds, the annotation interface, the variability-aware CRF-ID implementation, the alignment code, and the device CAD files will be made available upon publication.

## Author contributions

Z.Yu, H.L. and Y.Z. conceived the project. J.L., Z.Yu, Y.M., S.M. and W.Y. generated the transgenic strains. Z.Yu developed the microfluidic device and imaging protocol and acquired the hermaphrodite atlas and high-resolution whole-brain images; W.Y. acquired the male images; S.M. acquired functional recordings and performed neuron tracking with assistance from L.P.A. and W.A.C. Z.Yu developed the detection models, annotation and analysis tools, and coordinated the annotation and consensus process. Z.Yu, W.Y., Y.M., Z.Yang, H.J.L. M.G. and S.K. performed neuron annotations with Y.Z.’s supervision; Z.Yu, W.Y., Y.M. and J.L. established the annotation criteria. Z.Yu developed the computational identification, alignment and atlas construction methods, built the hermaphrodite and male atlases, and performed their validation and applications. Z.Yu, Z.Yang and W.Y. performed the anatomical and behavioral experiments. Z.Yu analyzed data and prepared the figures, with contributions from W.Y., Z.Yang, Y.M. and S.M. Z.Yu and H.L. drafted the main text and all authors contributed to the edits. H.L. and Y.Z. supervised the project and acquired funding.

## Acknowledgments

We thank Caenorhabditis Genetics Center for strains (funded by NIH Office of Research Infrastructure Programs P40 OD010440) for strains. H.L and Y.Z are funded by NIH (MH130064 and NS115484). H.L. is funded by NIH AG082039, NSF 1764406, and Simons Foundation 594594. We thank D.Buggs, A.Chakragiri, and A.Eaton for assistance with screening of generated transgenic strains.

## Supplementary information

**Supplementary Fig. 1:**
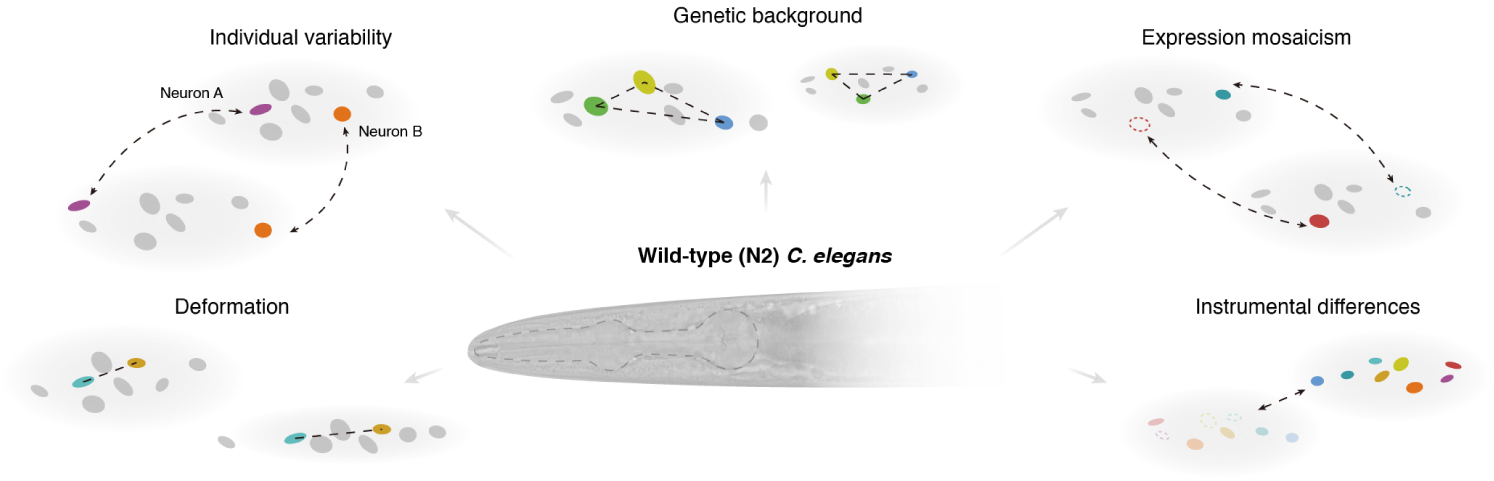
Discrepancy between a static atlas and a new dataset make accurate neural identification difficult. Several sources of discrepancy are possible for *C. elegans*, including positional variability between individuals, genetic background, expression mosaicism, device-induced deformation, and instrumentation differences.

**Supplementary Fig. 2:**
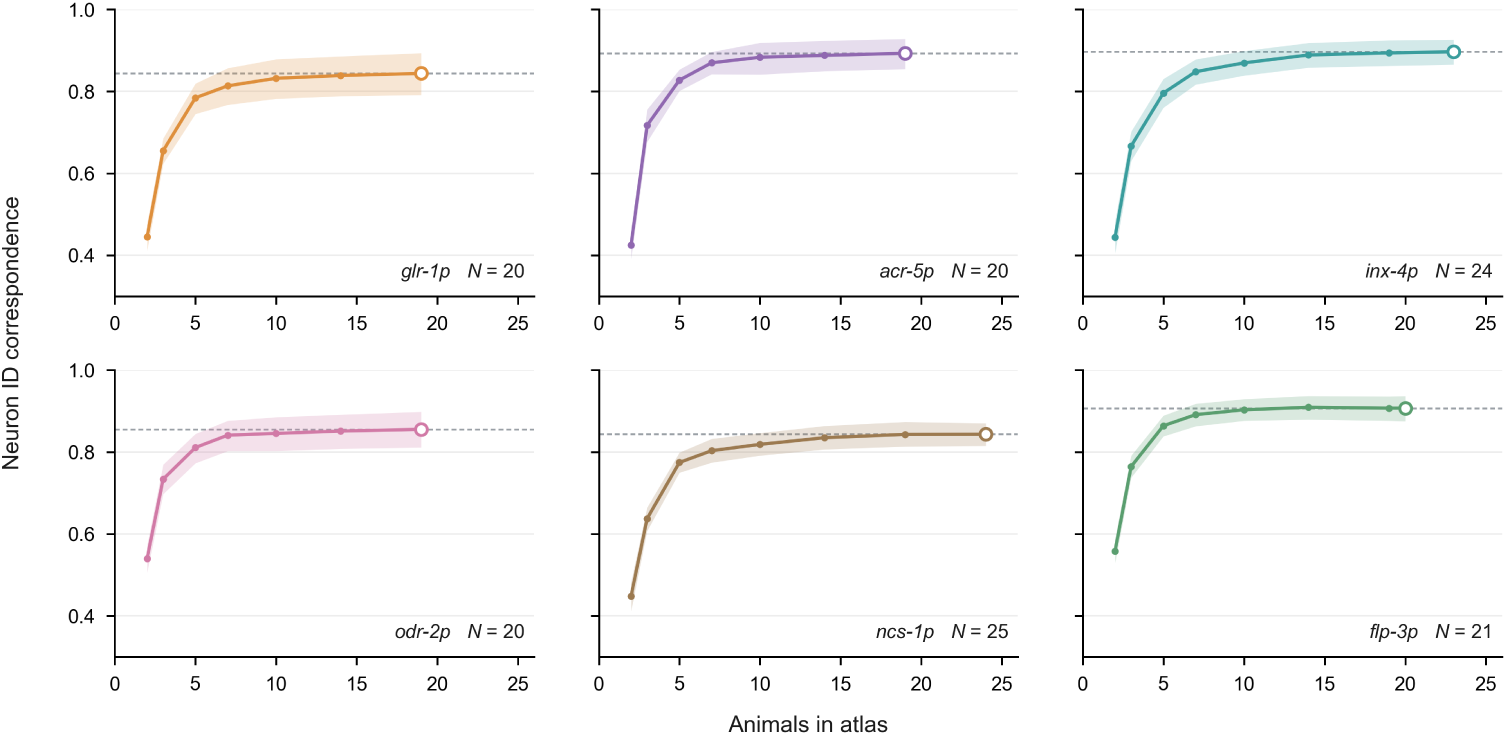
Identification accuracy versus number of worms used to build the atlas. The accuracy, shown as the neuron ID correspondence, plateaus beyond roughly ten worms per strain.

**Supplementary Fig. 3:**
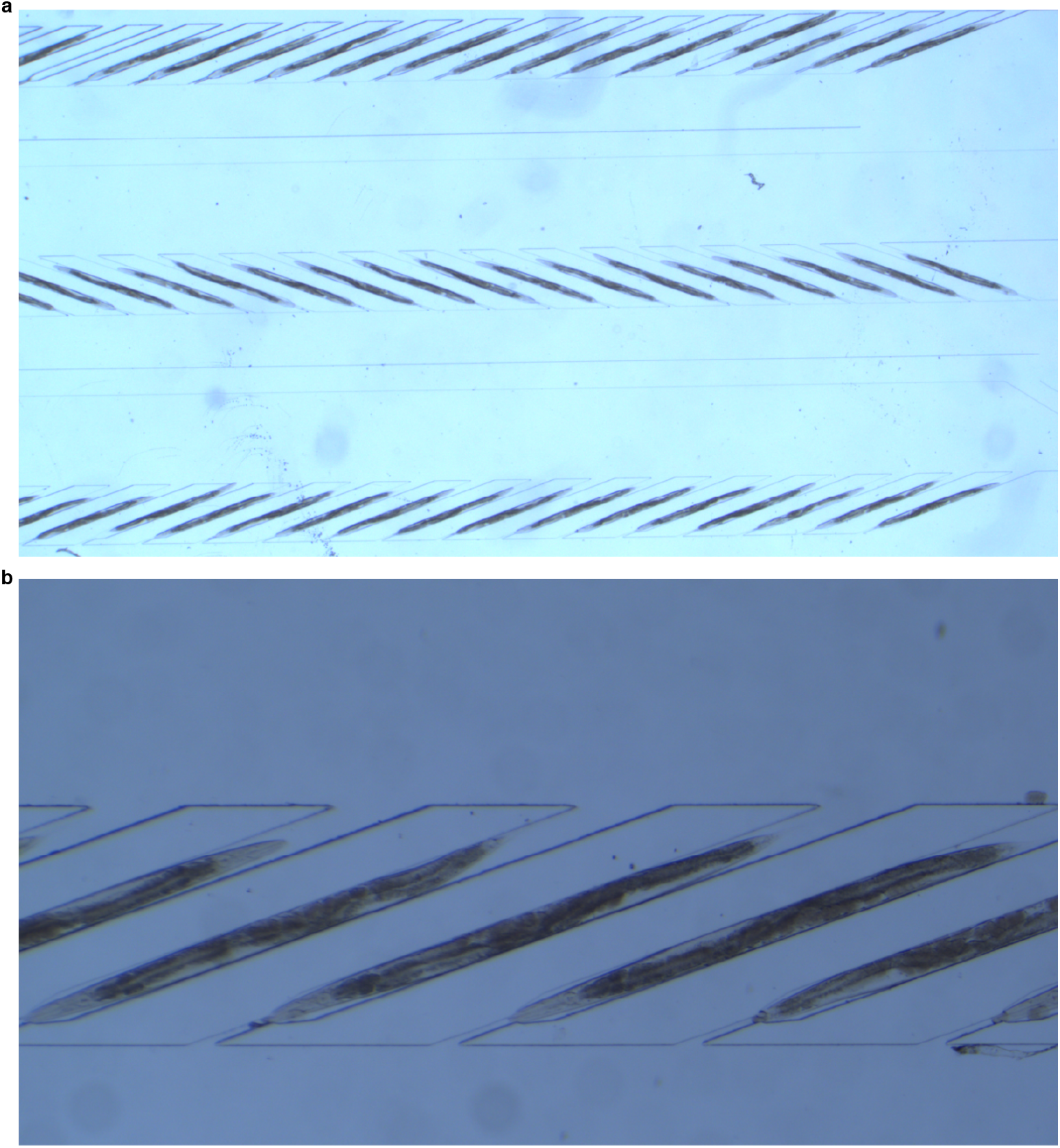
Array-style microfluidic device. **a**. 4× and **b**. 20× brightfield images of the microfluidic device loaded with worms. Each device can be loaded with approximately 100 worms in about five minutes of hands-on time.

**Supplementary Fig. 4:**
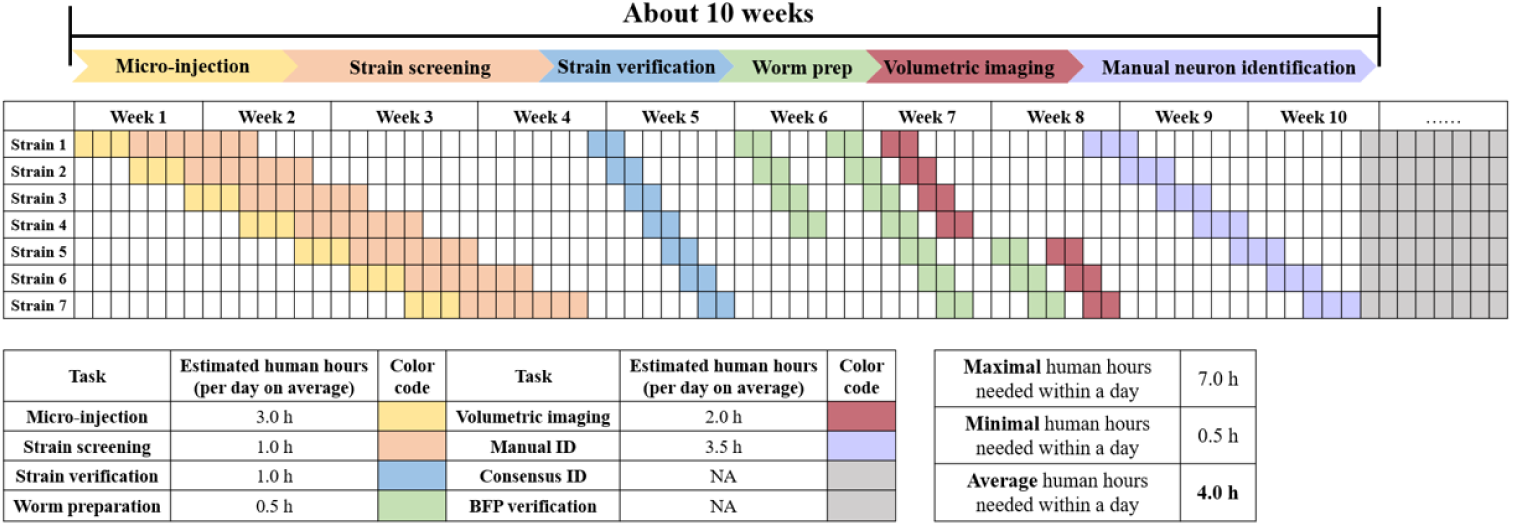
Estimated timeline and human labor for building an atlas. (Top) Building an atlas of seven strains is expected to take approximately ten weeks when the work is done in parallel. (Bottom) Estimated per-stage hands-on hours.

**Supplementary Fig. 5:**
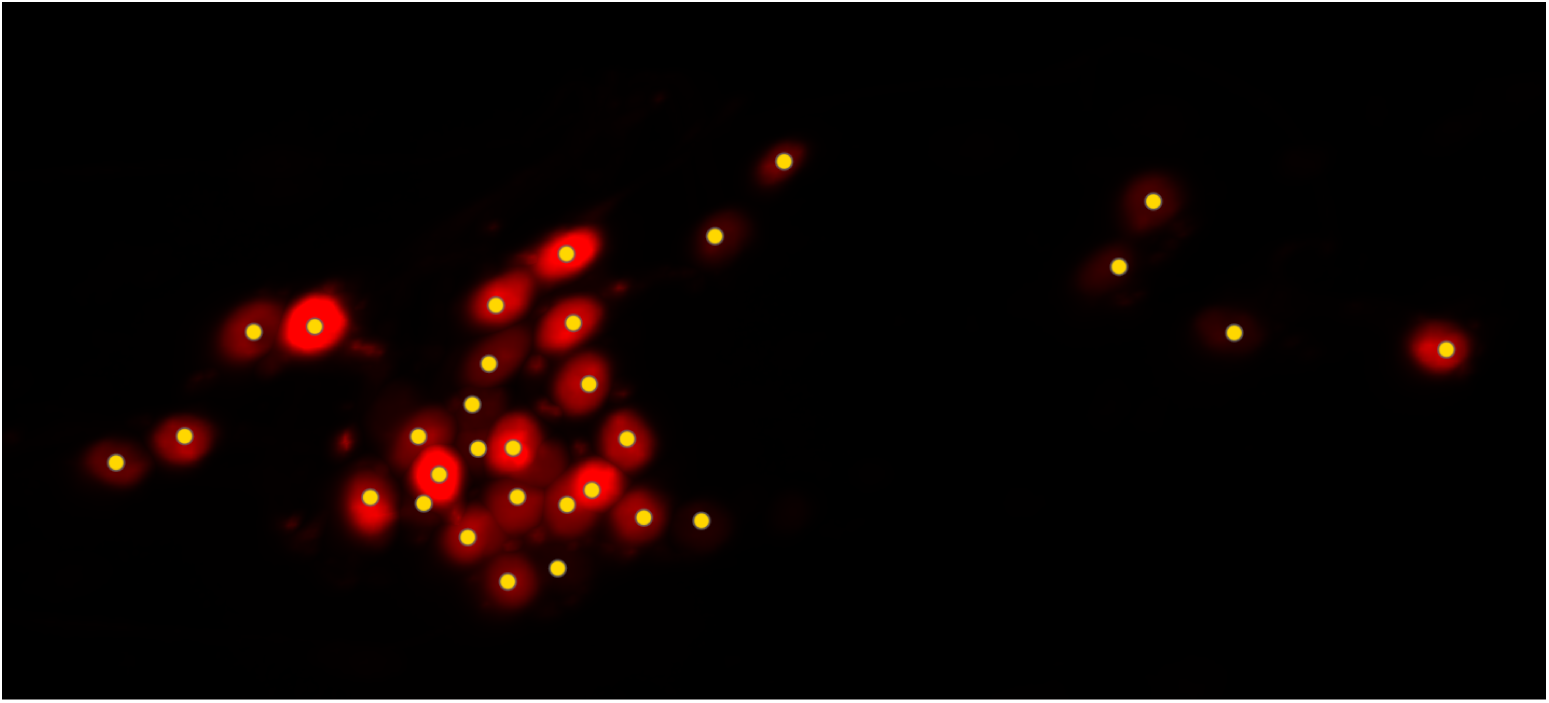
Automatic detection of labeled nuclei in the mCherry channel. Example of a multi-cell worm with corresponding neuron detections.

**Supplementary Fig. 6:**
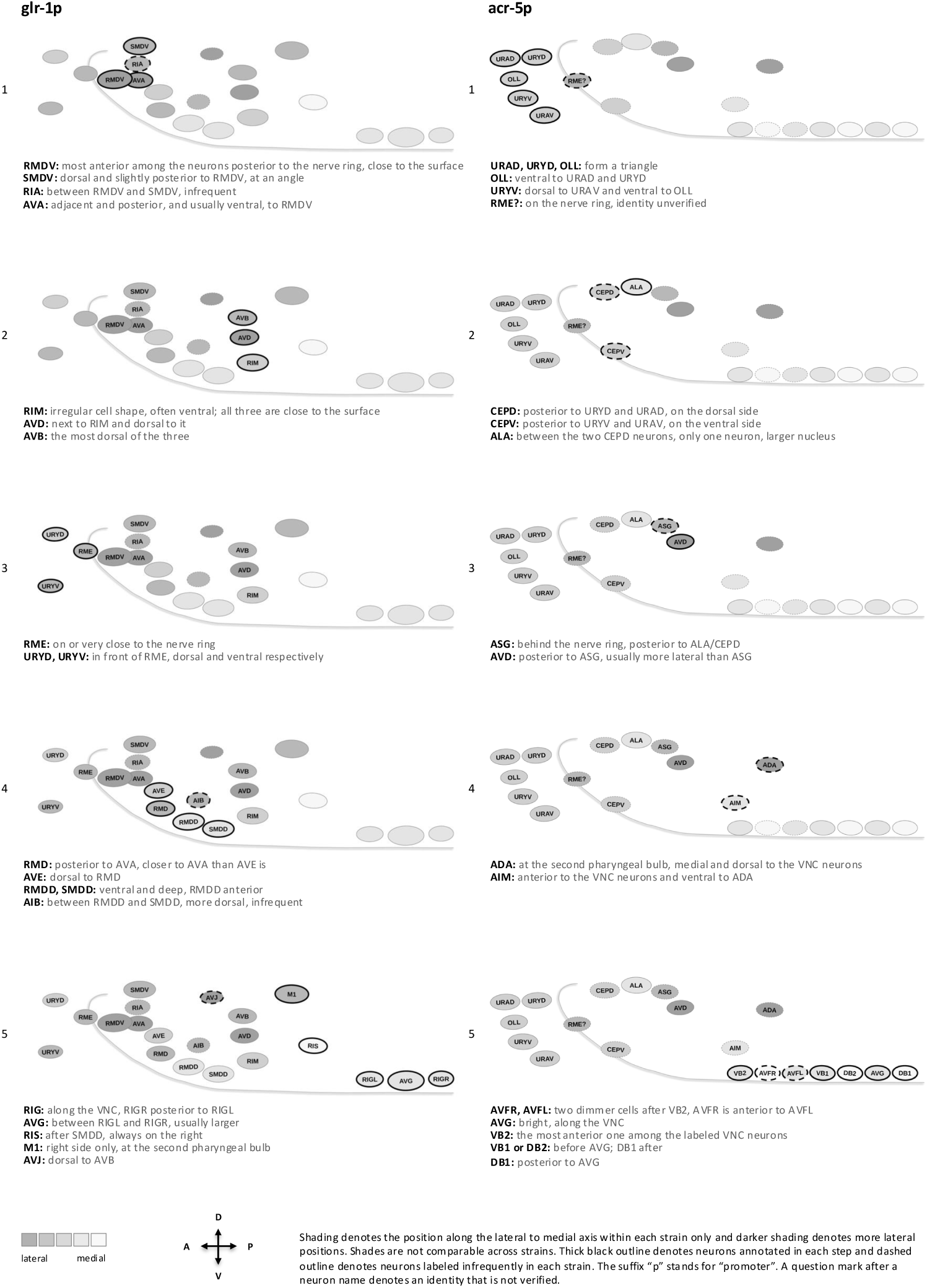
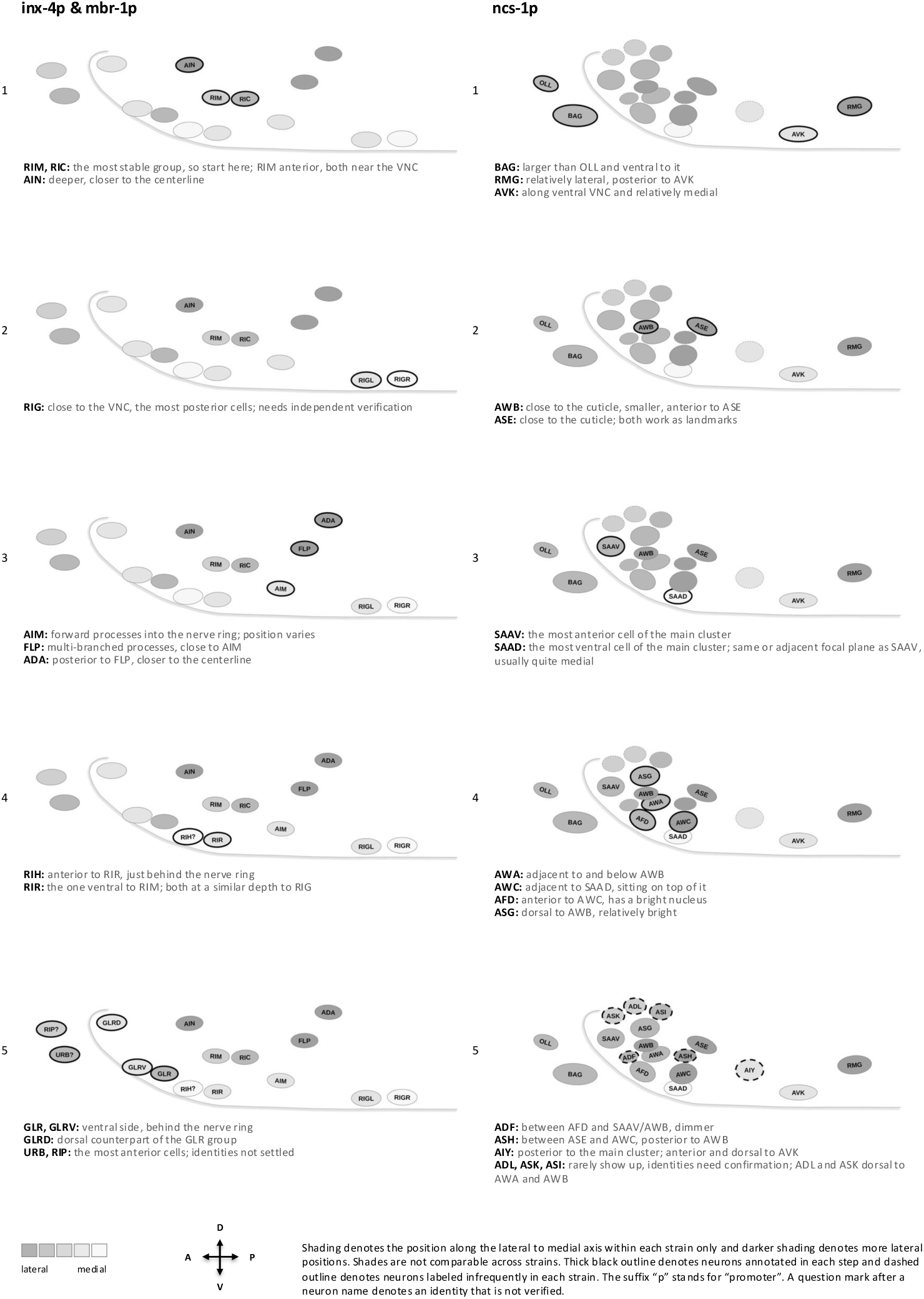
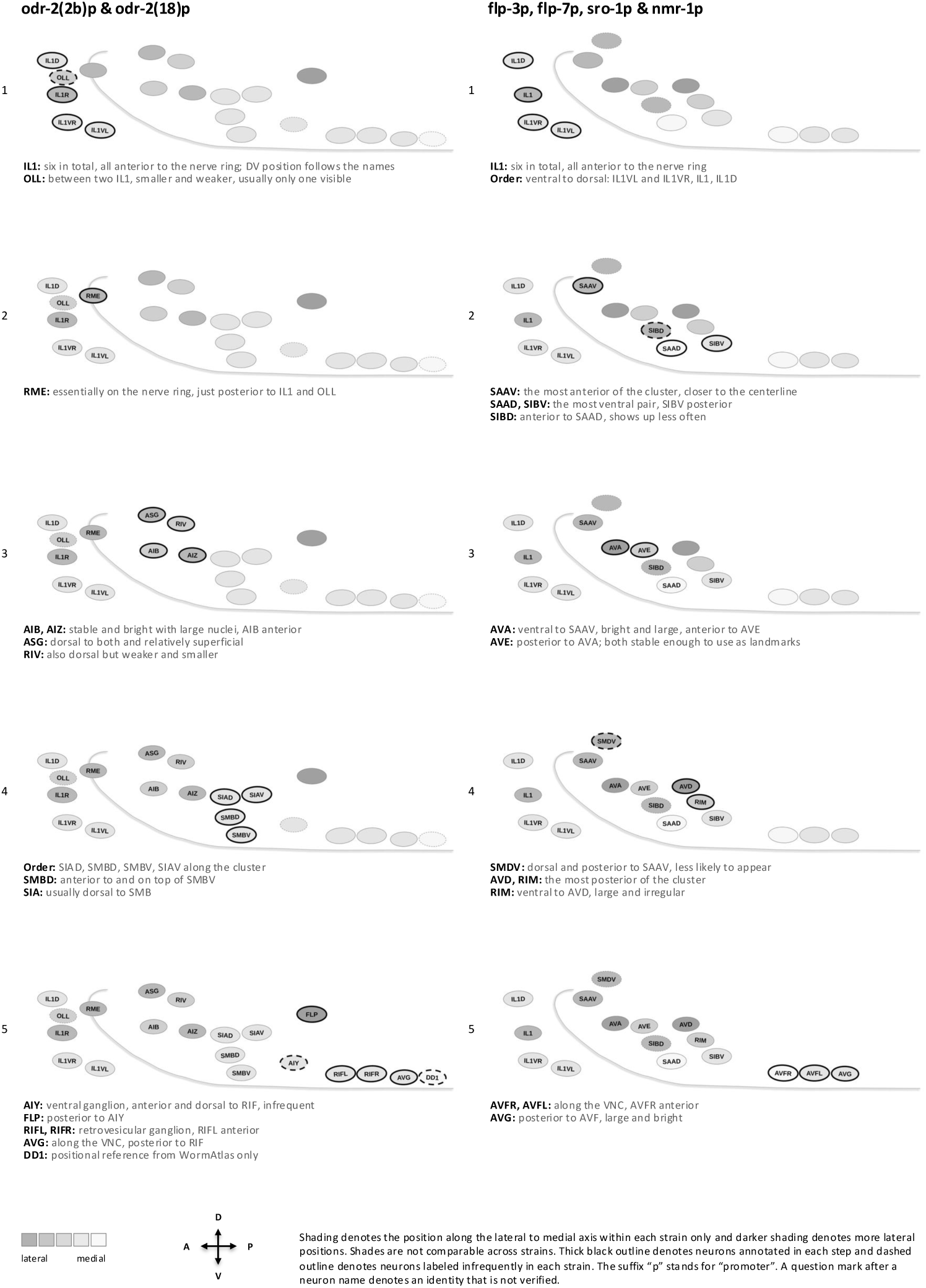
Annotation criteria for *glr-1p*, *acr-5p*, *inx-4p & mbr-1p*, *ncs-1p*, *odr-2(2b)p & odr-2(18)p*, and *flp-3p, flp-7p, sro-1p & nmr-1p*. Shading denotes the position along the lateral to medial axis within each strain only and darker shading denotes more lateral positions. Shades are not comparable across strains. Thick black outline denotes neurons annotated in each step and dashed outline denotes neurons labeled infrequently in each strain. The suffix “p” stands for “promoter”. A question mark after a neuron name denotes an identity that is not verified.

**Supplementary Fig. 7:**
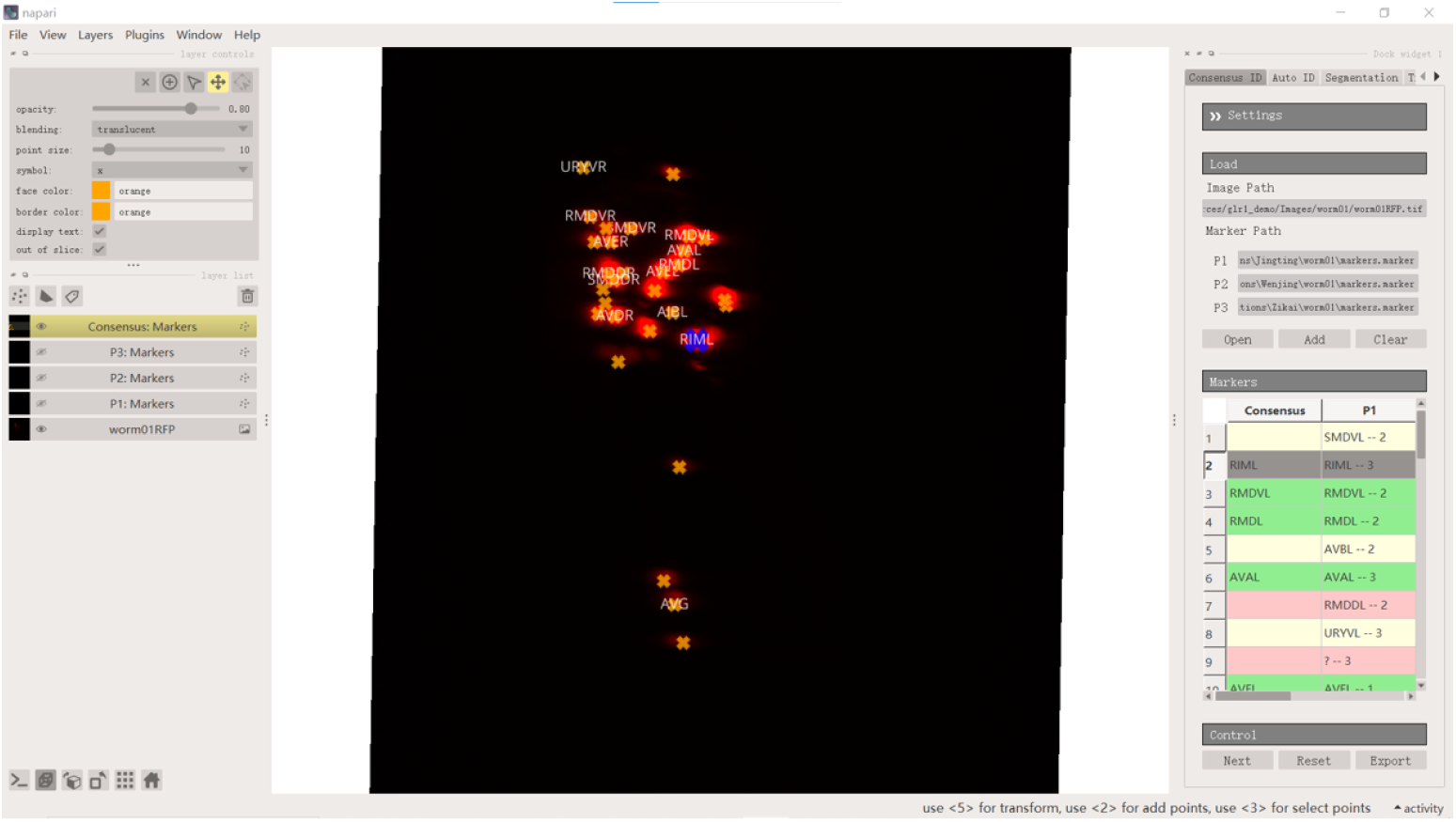
Annotation interface. napari-based interface showing detected nuclei with consensus labels overlaid (left) and, for each nucleus, the labels and confidence scores of each annotator alongside the consensus (right).

**Supplementary Fig. 8:**
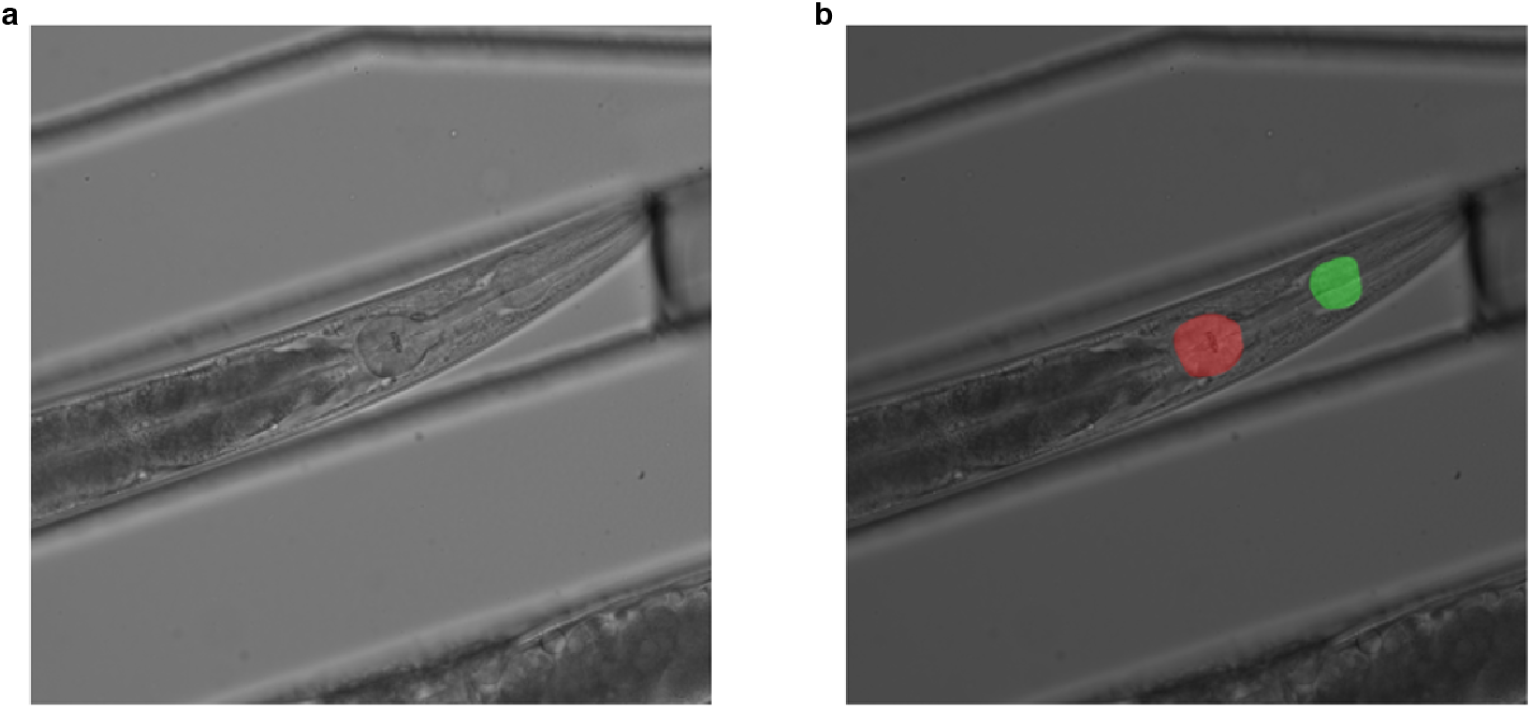
Automatic bulb detection in brightfield images using a Vision Transformer (ViT) model. **a**. A representative brightfield image in the test set and **b**. the ViT mask prediction for the two bulbs. The center of the bulbs are estimated from the masks.

**Supplementary Fig. 9:**
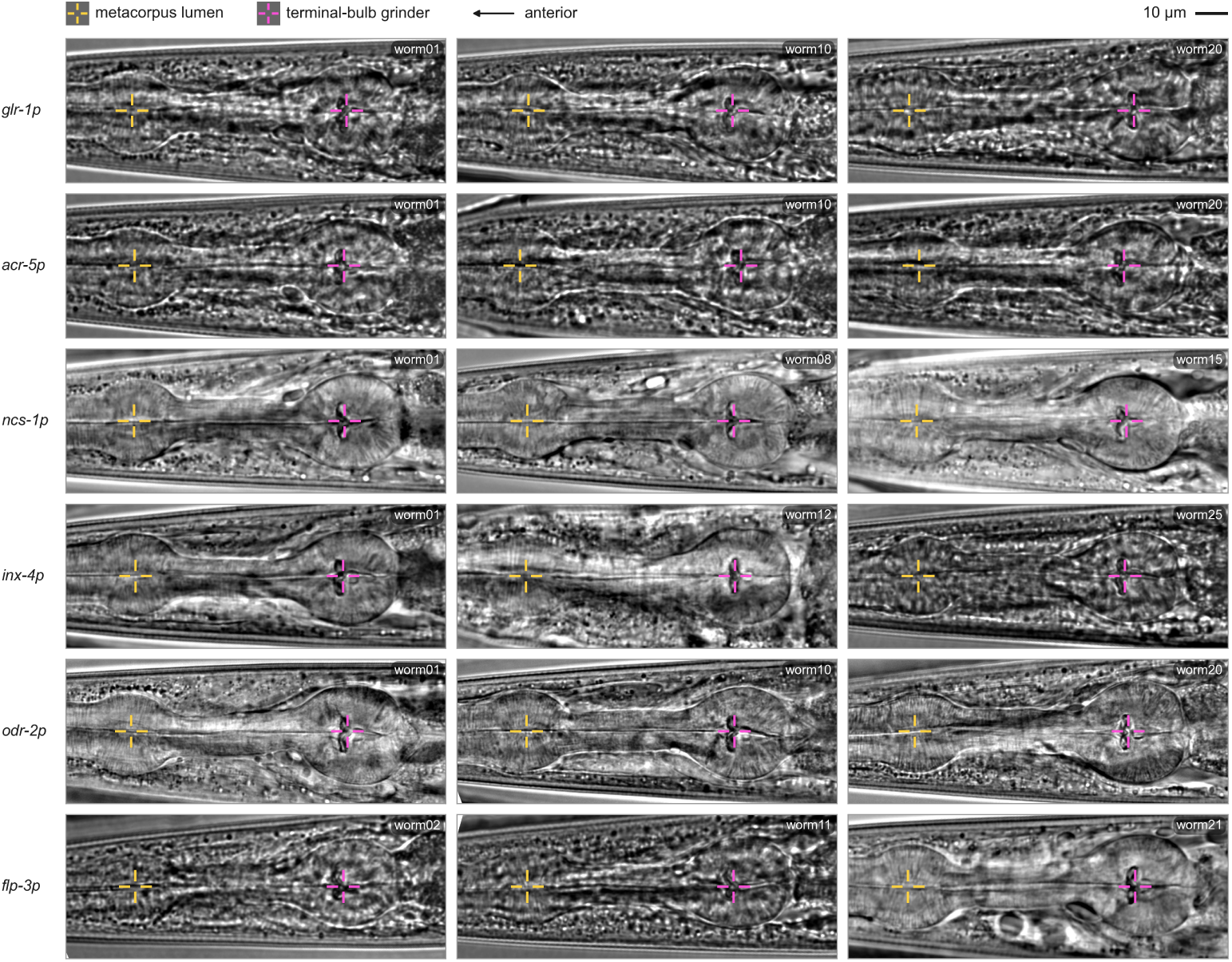
Terminal bulb detection in brightfield images. Orange and magenta markers indicate the detected first and second bulbs, respectively. The origin and AP axis will be calculated based on the detected centers.

**Supplementary Fig. 10:**
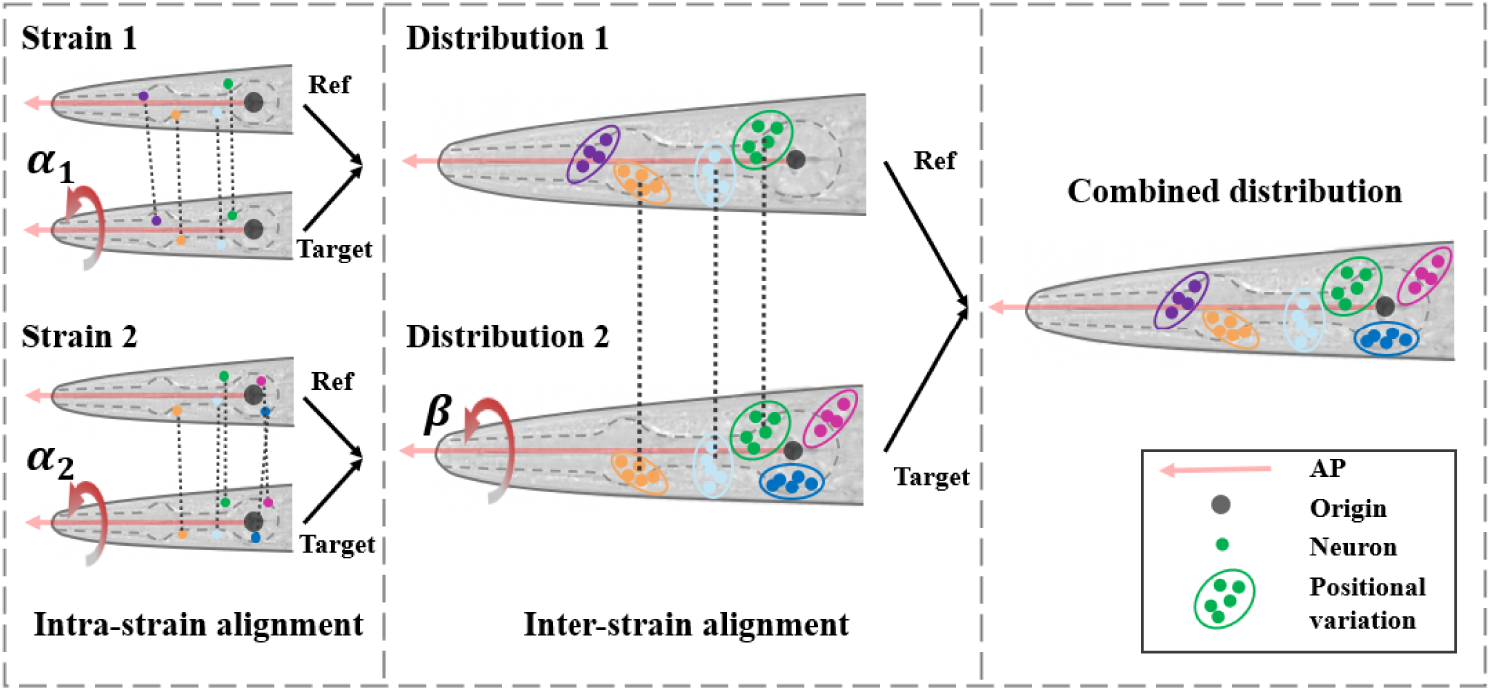
Two-stage anatomy-assisted alignment. Intra-strain rotation about the AP axis, followed by the inter-strain alignment at the level of positional distributions.

**Supplementary Fig. 11:**
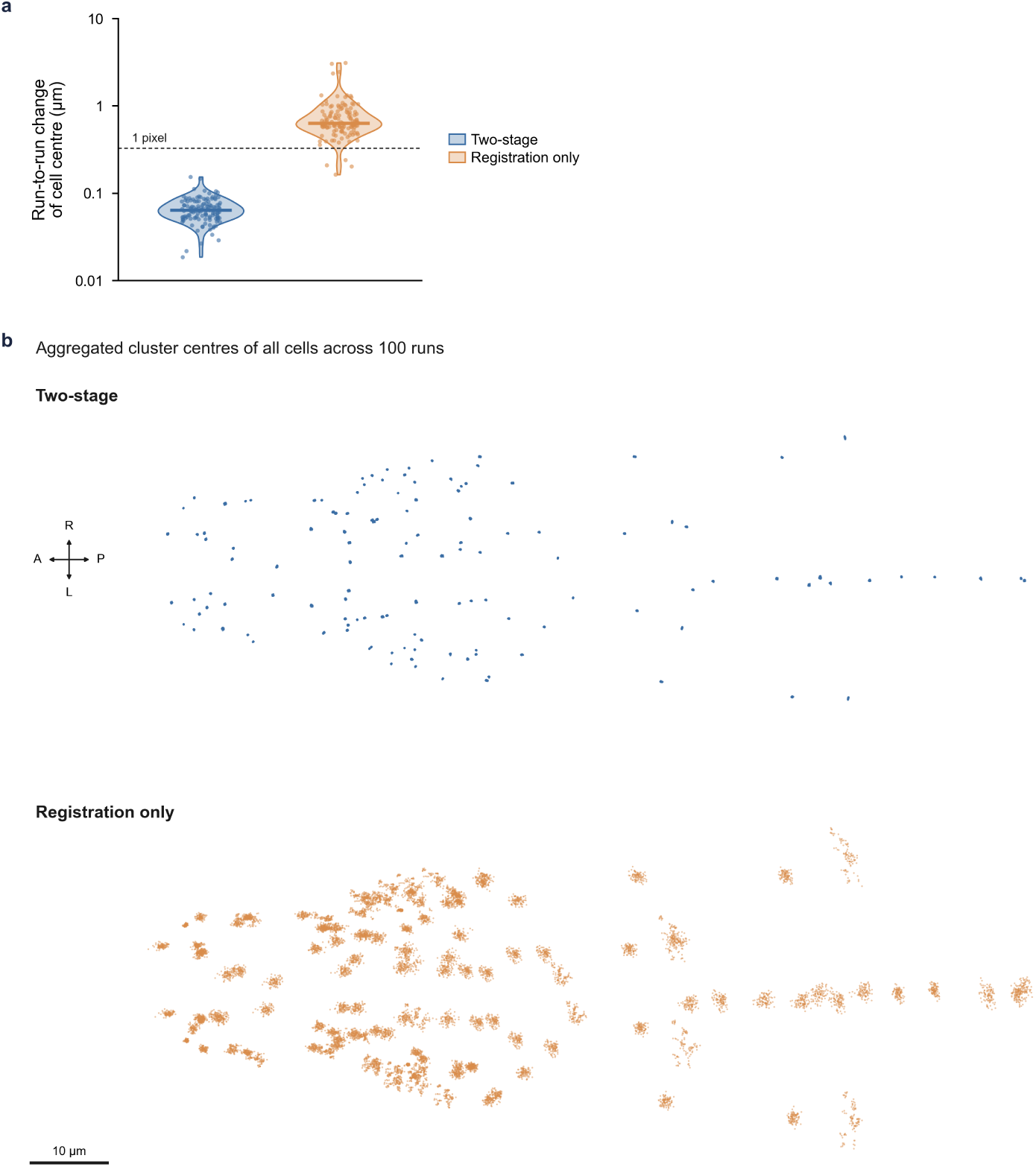
Two-stage alignment versus registration-only alignment. **a**. The run-to-run difference of the resulting atlases starting from different reference worms for alignment. Among the 100 random atlas-building runs, the proposed two-stage methods exhibit median cluster center drifting below 0.1 *µ*m (the voxel size is 0.3 *µ*m), which is substantially more stable than using a registration-based method. **b**. The aggregated cluster center distribution across 100 independent runs for both methods, displayed using the same scale bar. The two-stage method has much tighter spread compared to the registration based method.

**Supplementary Fig. 12:**
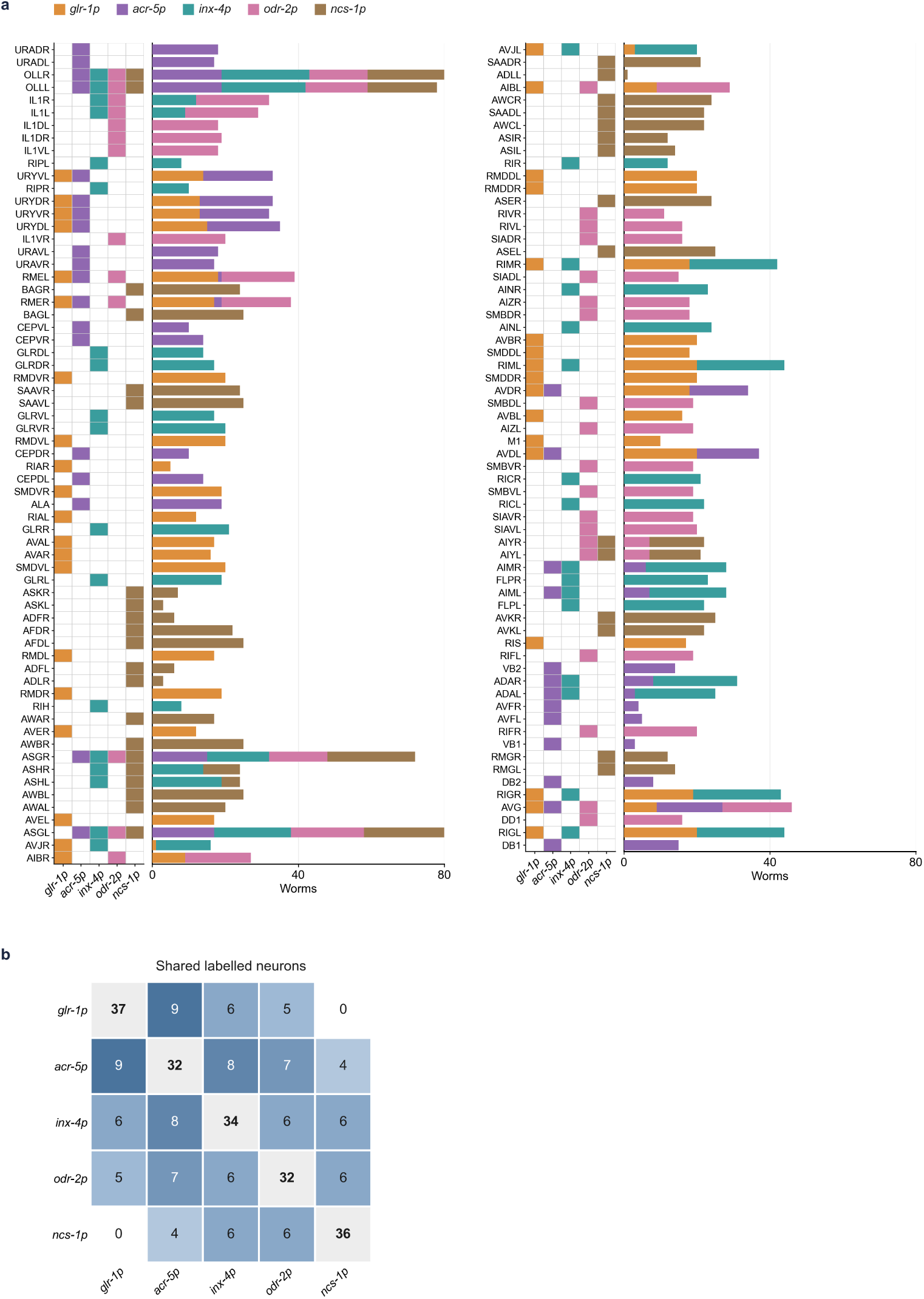
Coverage of the atlas strains at the level of individual neurons. **a**. Neurons labeled in each of the five atlas-building strains and the number of worms in which each neuron was expressed (stacked by strain). Neurons are ordered from anterior to posterior. **b**. The overlapping among the 5 atlas-building strains.

**Supplementary Fig. 13:**
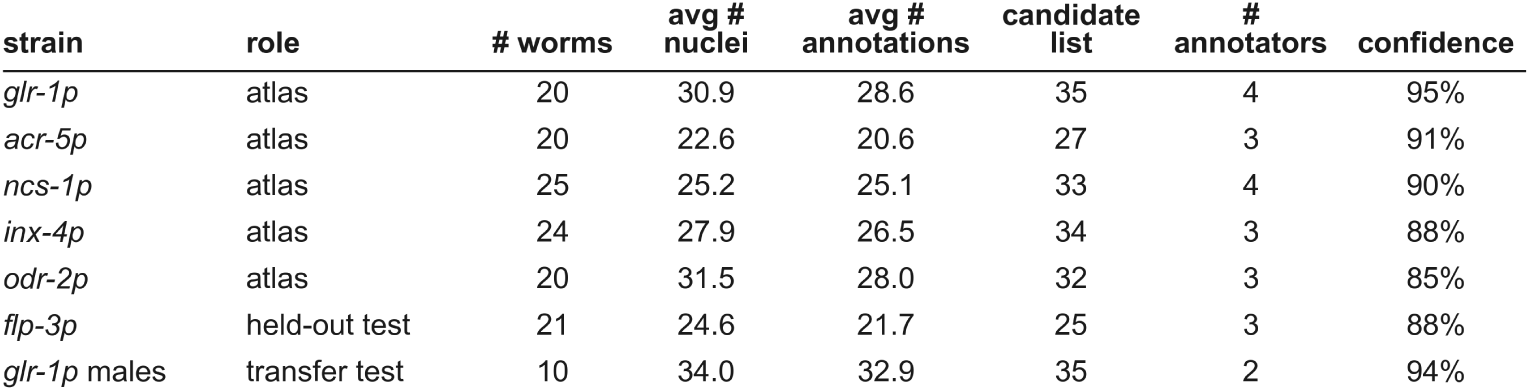
Summary of multi-cell datasets involved in this study.

**Supplementary Fig. 14:**
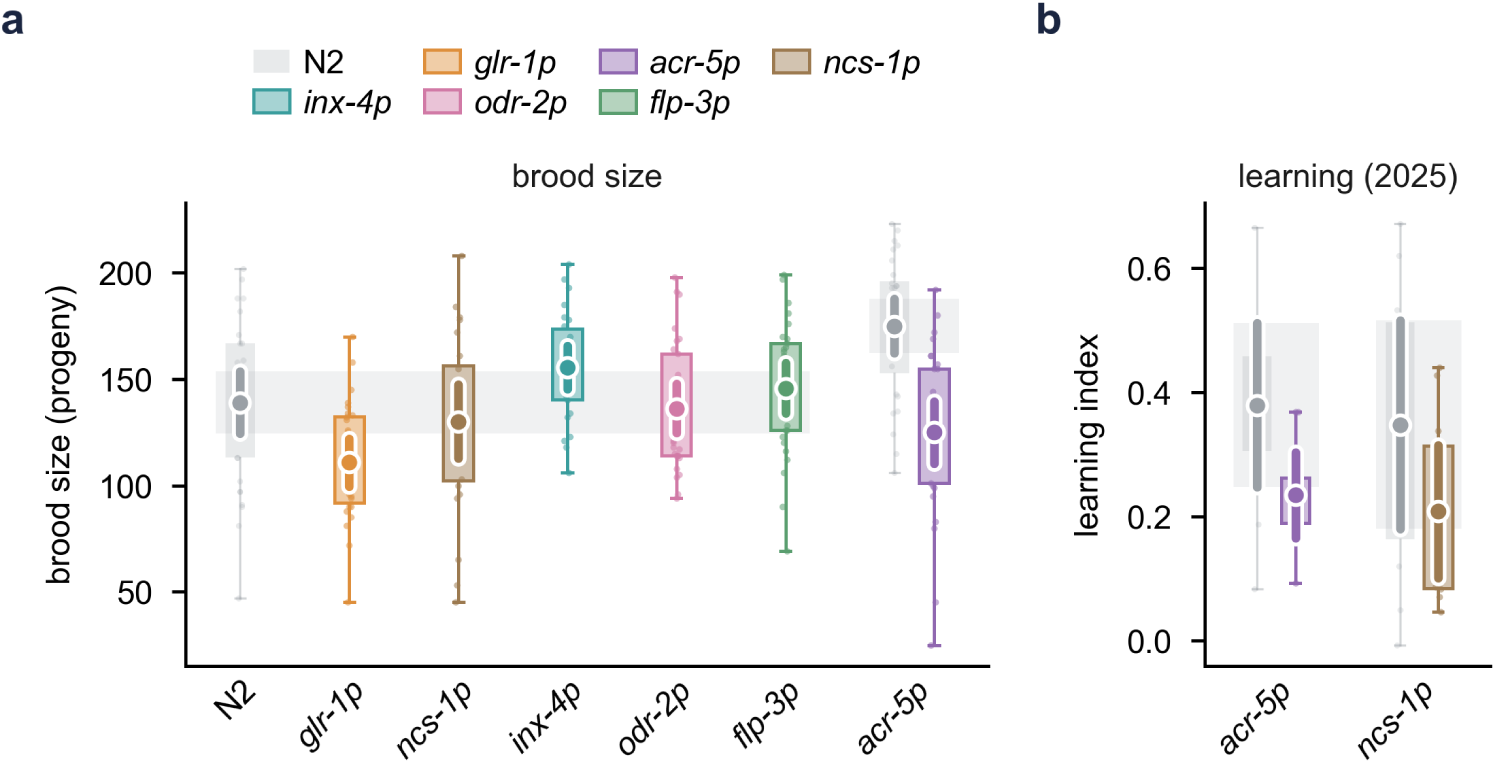
Additional assays for strain health. **a**. Brood size of the six different strains compared with N2. **b**. The additional learning assays for the *acr-5* and *ncs-1* strain. The learning data in Fig. 2c were collected after those shown in **b**, once the assay protocol had been refined; we include the earlier series here for transparency.

**Supplementary Fig. 15:**
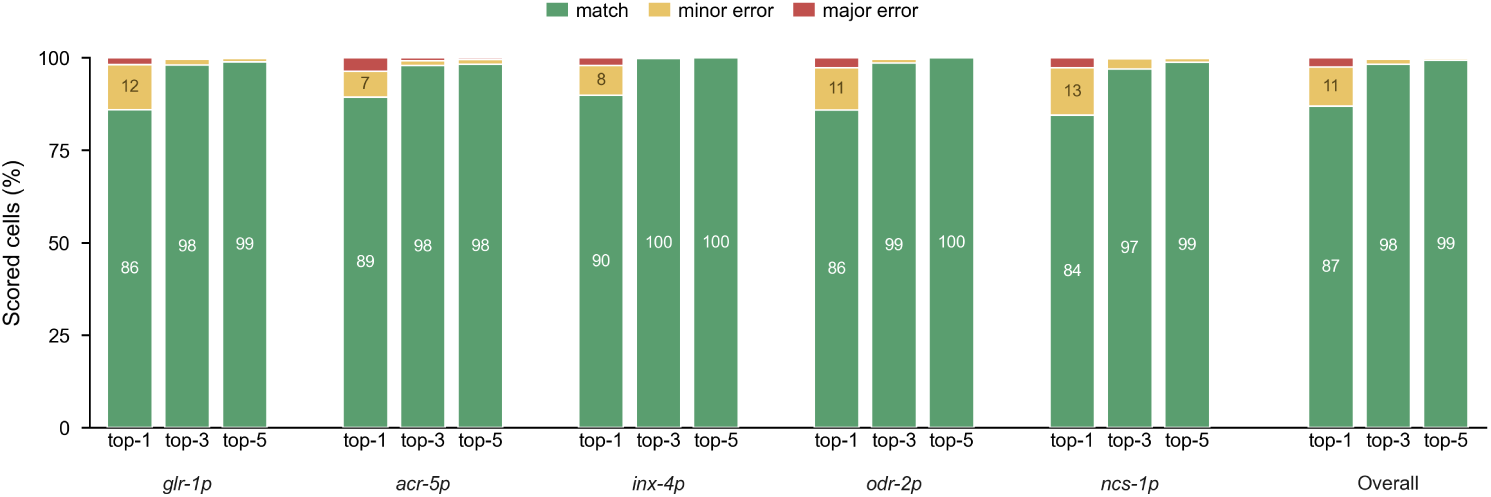
Error severity in strain-matched leave-one-out identification. Match (green), minor error (anatomical neighbor; yellow), and major error (red) for top-1, top-3, and top-5 predictions.

**Supplementary Fig. 16:**
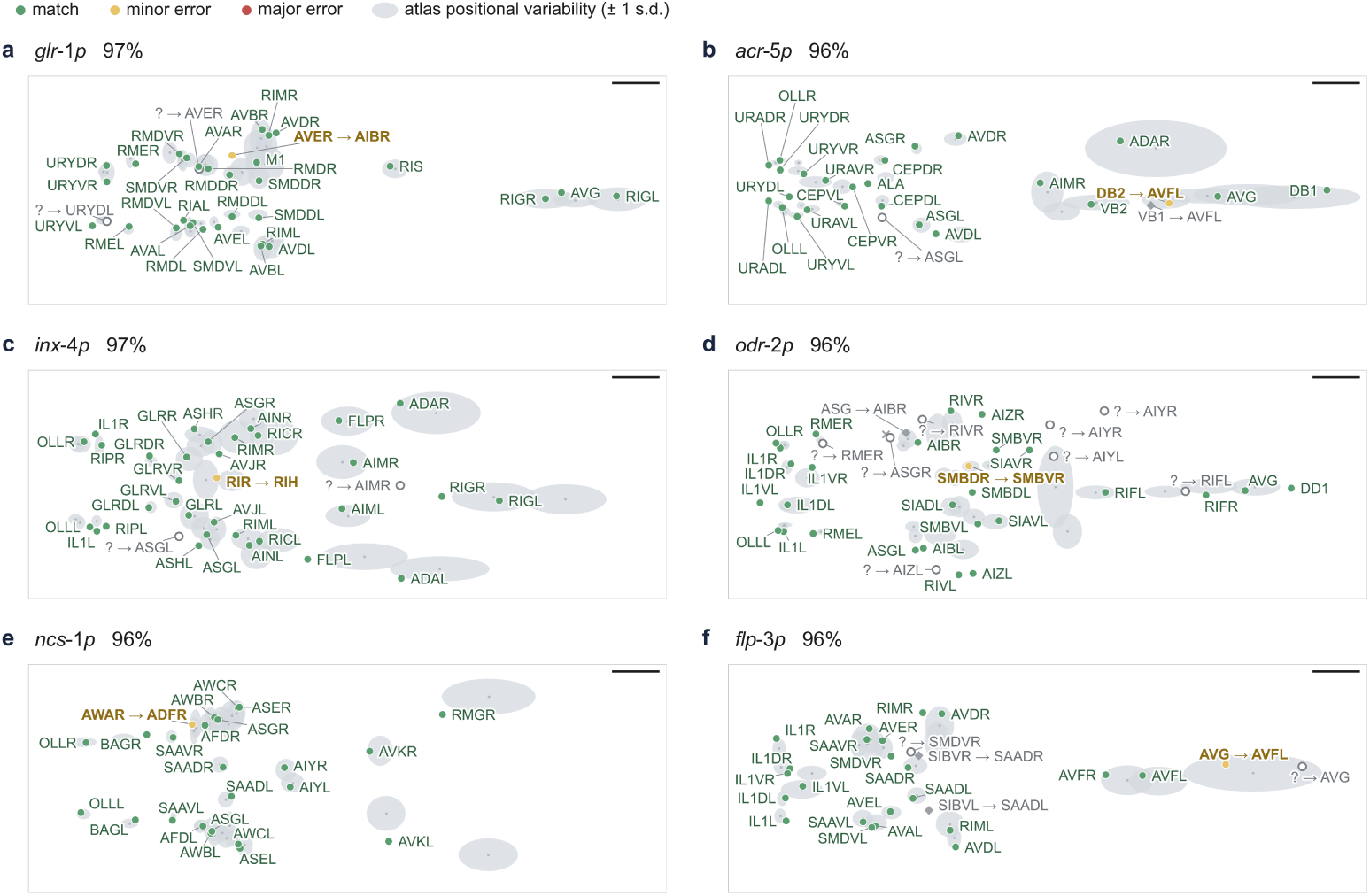
Representative examples of using strain-specific atlases to infer multi-cell data. Match (green), minor error (anatomical neighbor; yellow), and major error (red) for top-1, top-3 and top-5 predictions.

**Supplementary Fig. 17:**
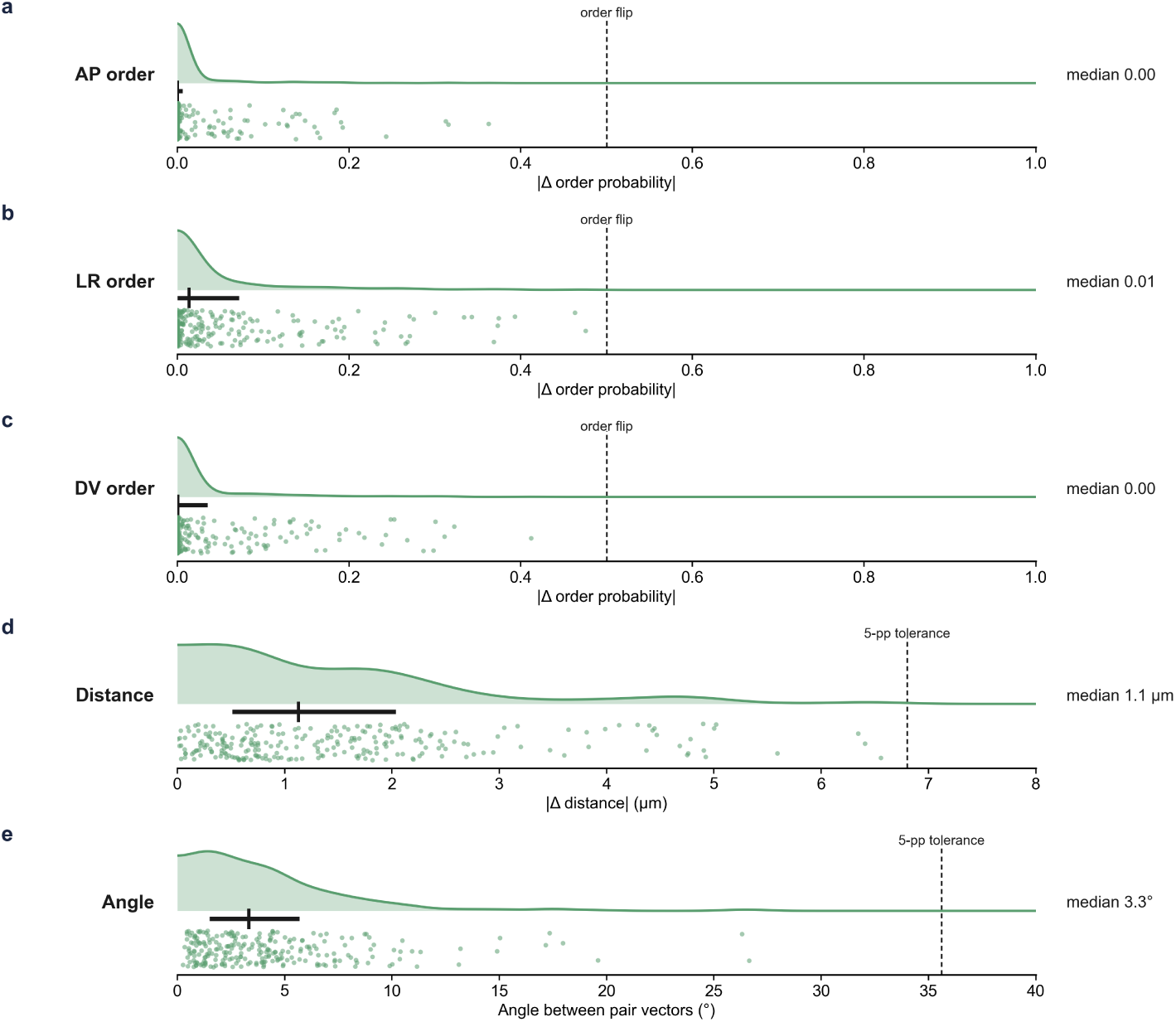
Differences between the brain-wide atlas and the *flp-3* atlas across all pairwise features considered in this work. From **a** to **e**, we compare the relative ordering along the AP, LR, DV axis, the pairwise distance, and the angles between corresponding neuron pairs respectively. Dots represent a specific neuron pair co-expressed in the *flp-3* strain; vertical lines represent median of the fitted distribution.

**Supplementary Fig. 18:**
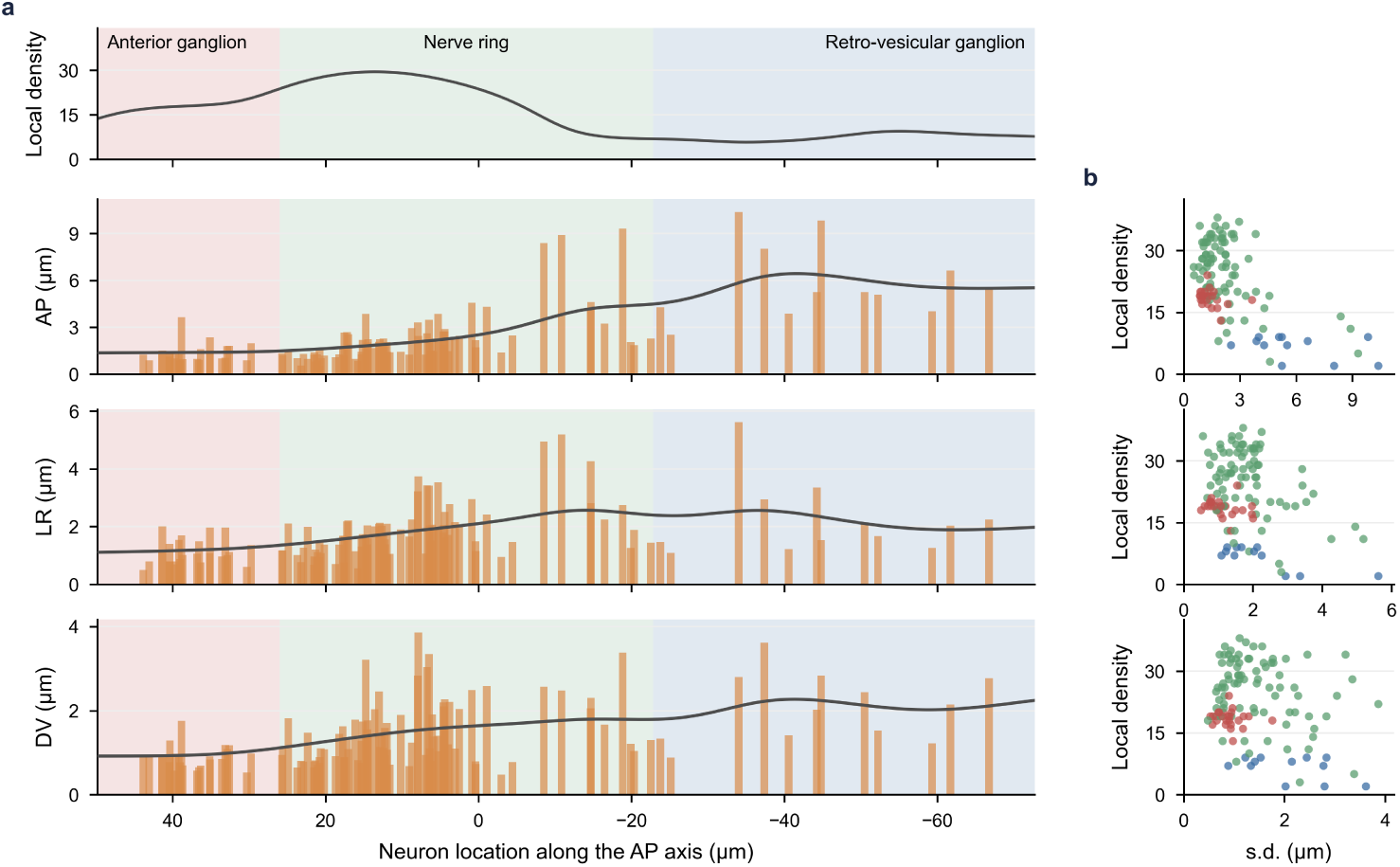
Positional variability along each anatomical axis. **a**. Local density versus AP/LR/DV variability. **b**. The correlation between local density and variability, with neuron classes colored by brain region.

**Supplementary Fig. 19:**
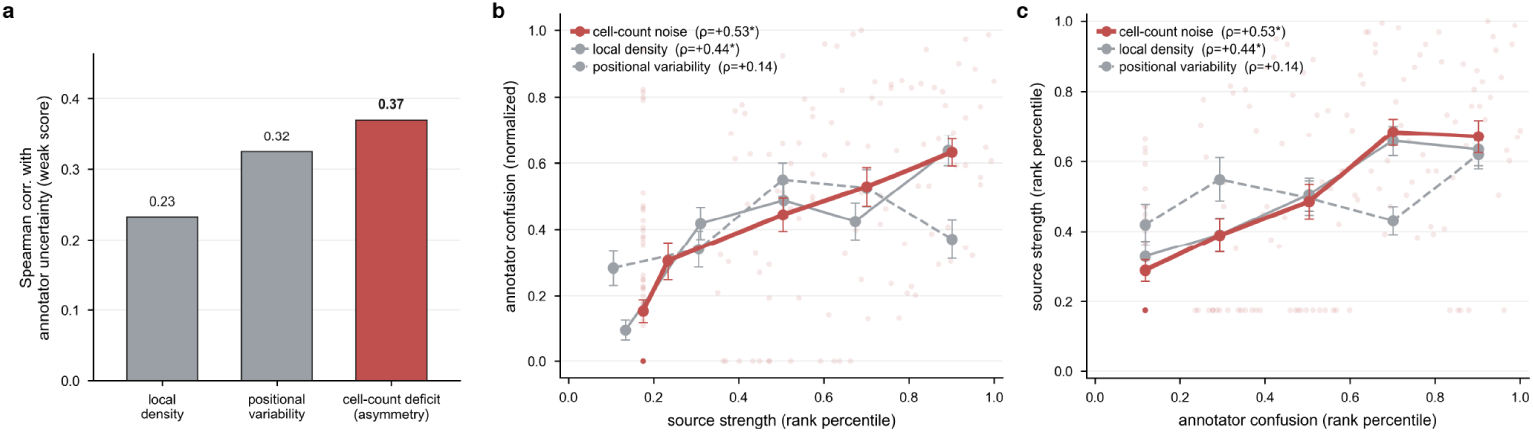
Sources of annotator confusion. **a**. Spearman correlation of annotator uncertainty with local density, positional variability, and cell-count deficit. **b and c**. Confusion against each source, shown with **b**. source strength and **c**. confusion on the x-axis.

**Supplementary Fig. 20:**
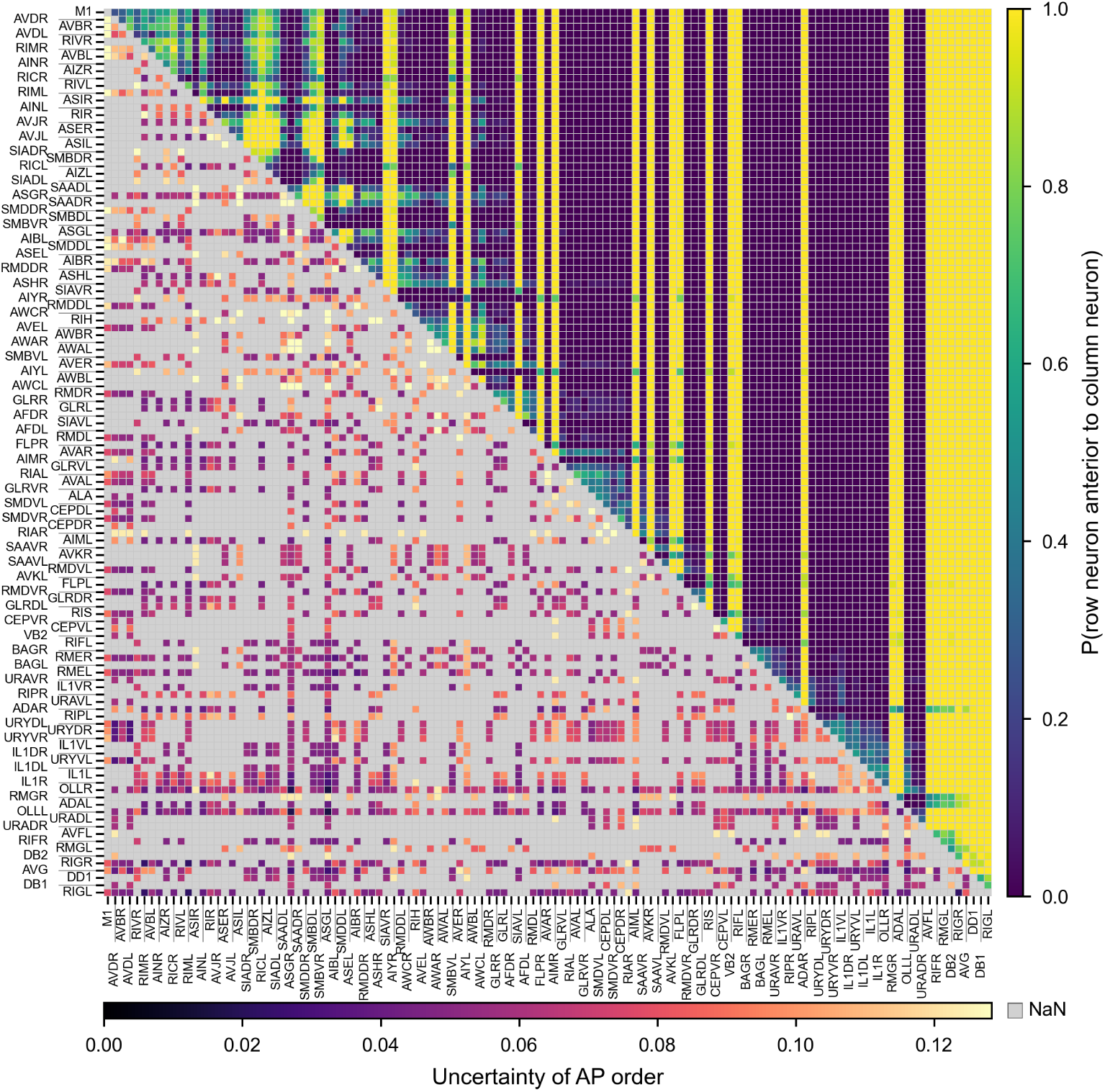
Relative ordering along the AP axis with embedded variability.

**Supplementary Fig. 21:**
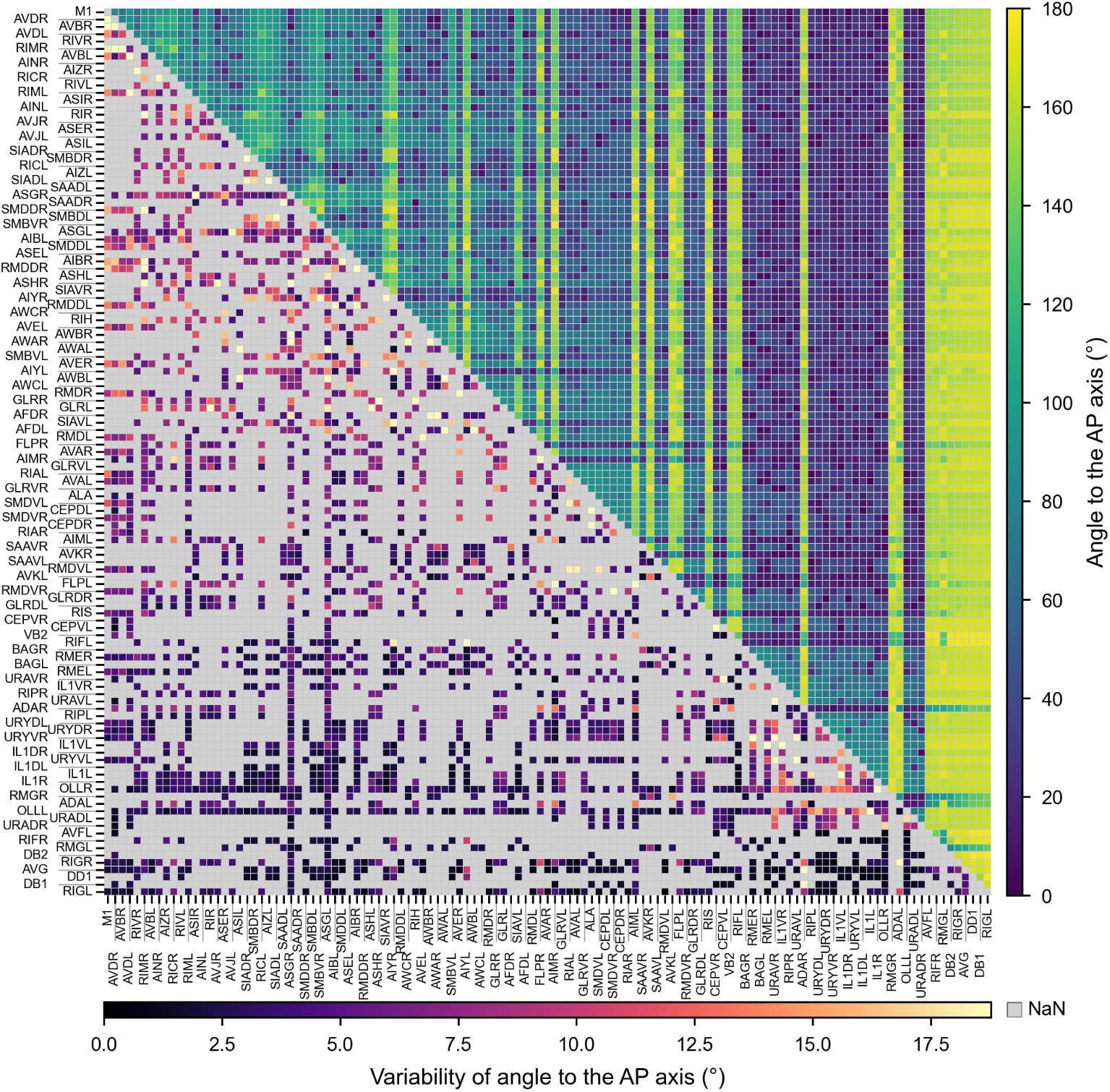
The angle between each neuron pair to the AP axis with embedded variability.

**Supplementary Fig. 22:**
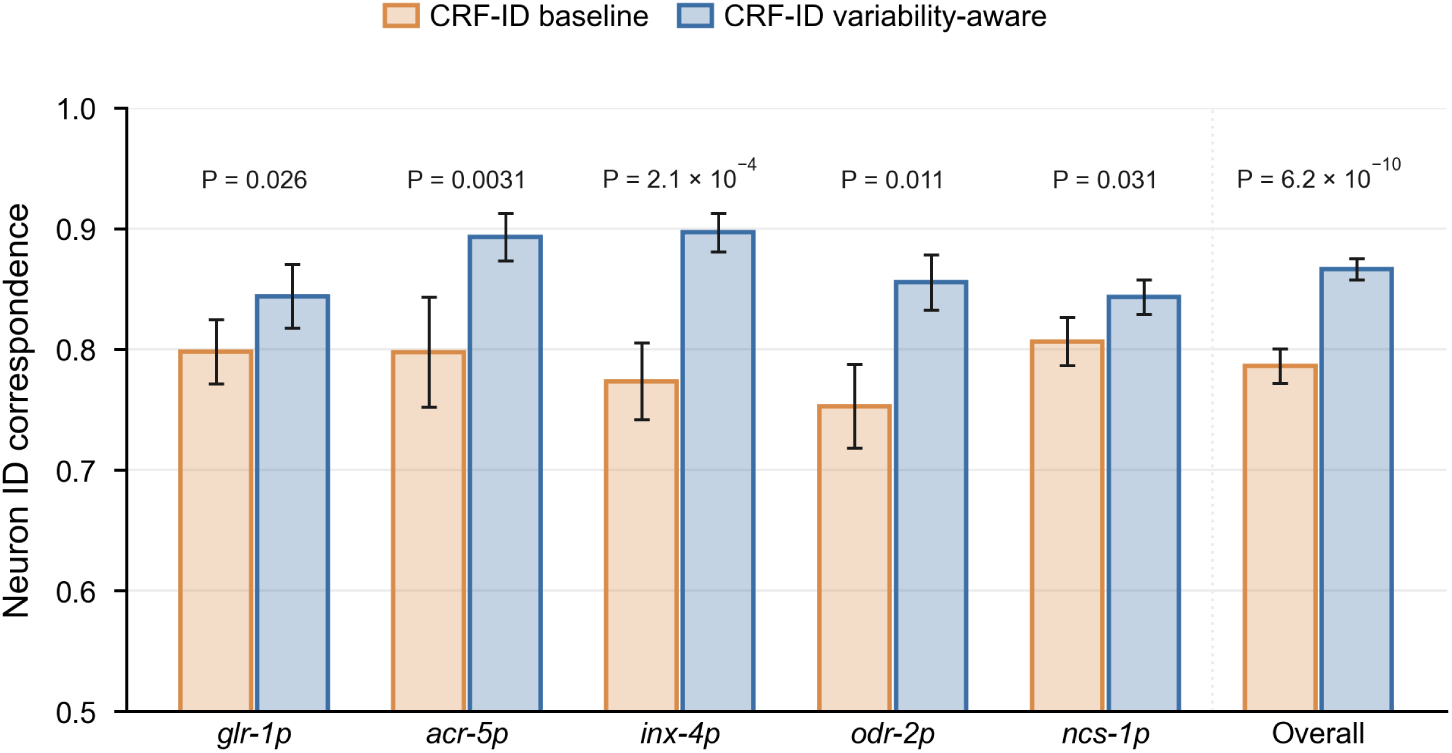
The comparison between the baseline CRF-ID and the variability-aware CRF-ID across all multi-cell strains. Error bars represent SEM.

**Supplementary Fig. 23:**
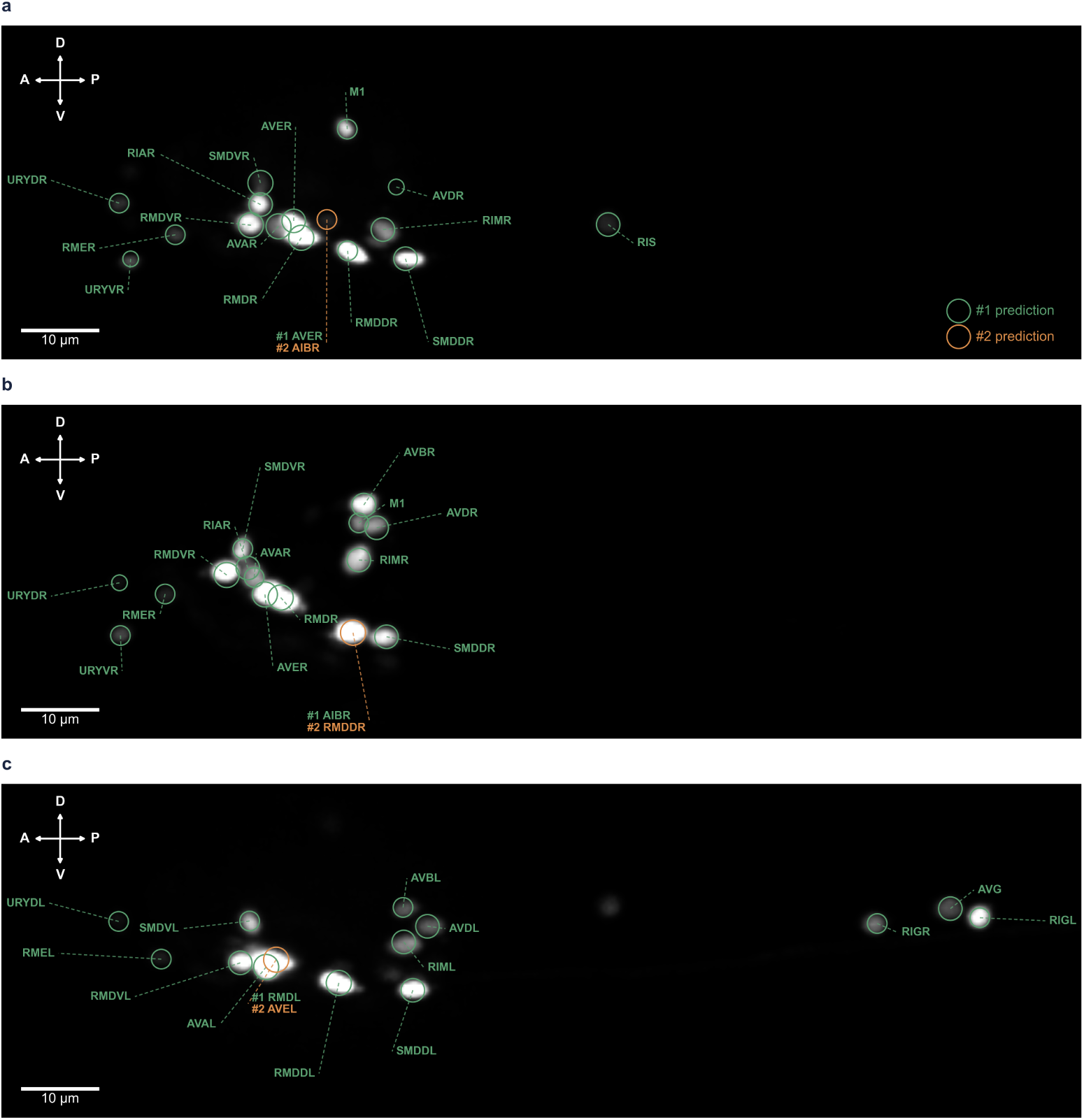
More examples of CRF-ID predictions using the hermaphrodite atlas to infer male worms. Green, rank-1 prediction; orange, rank-2 prediction.

**Supplementary Fig. 24:**
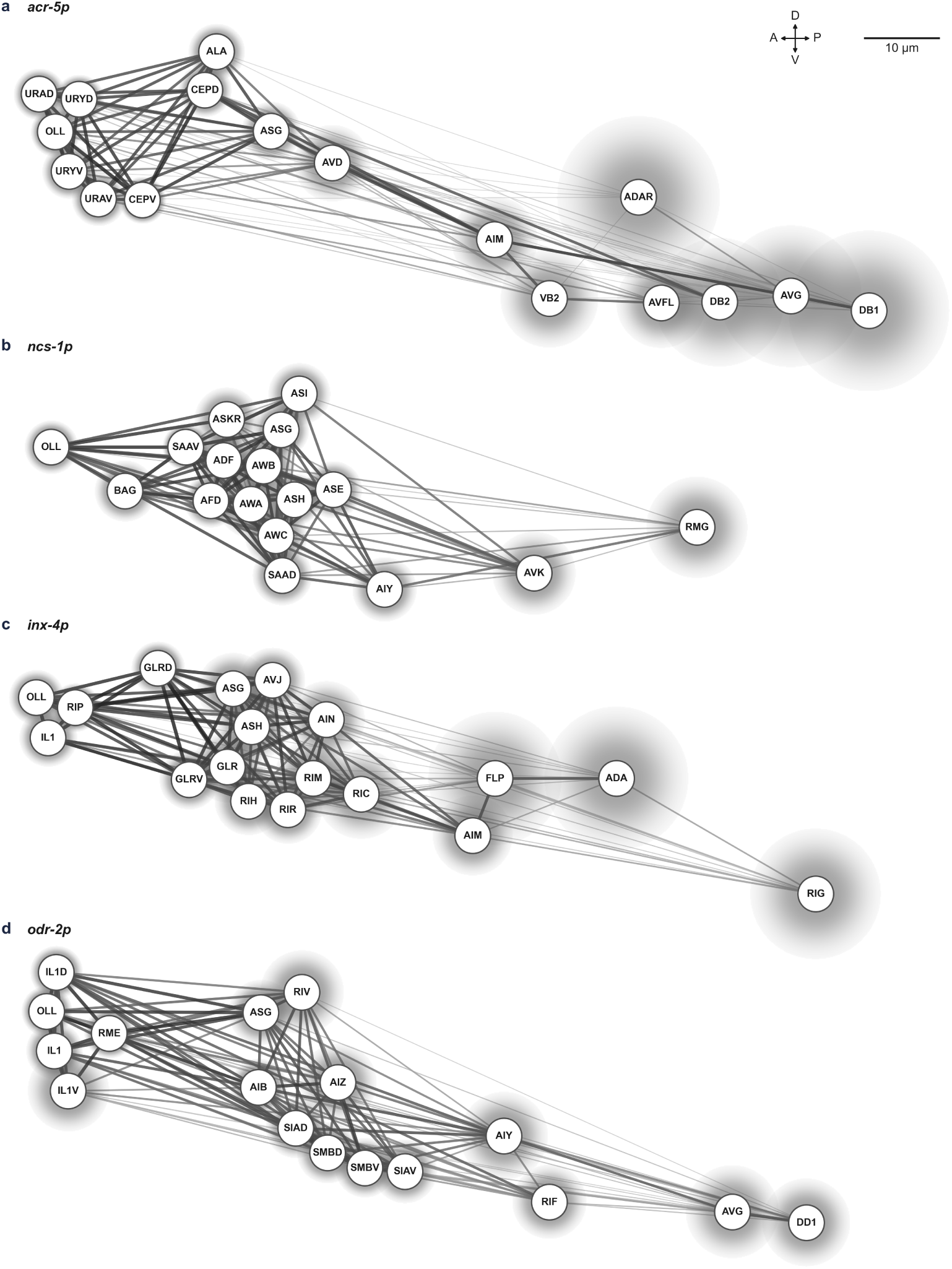
Variability atlas for other multi-cell strains. Relational graph of the **a**. *acr-5p*, **b**. *ncs-1p*, **c**. *inx-4p*, and **d**. *odr-2p* neurons in the variability-aware atlas.

**Supplementary Fig. 25:**
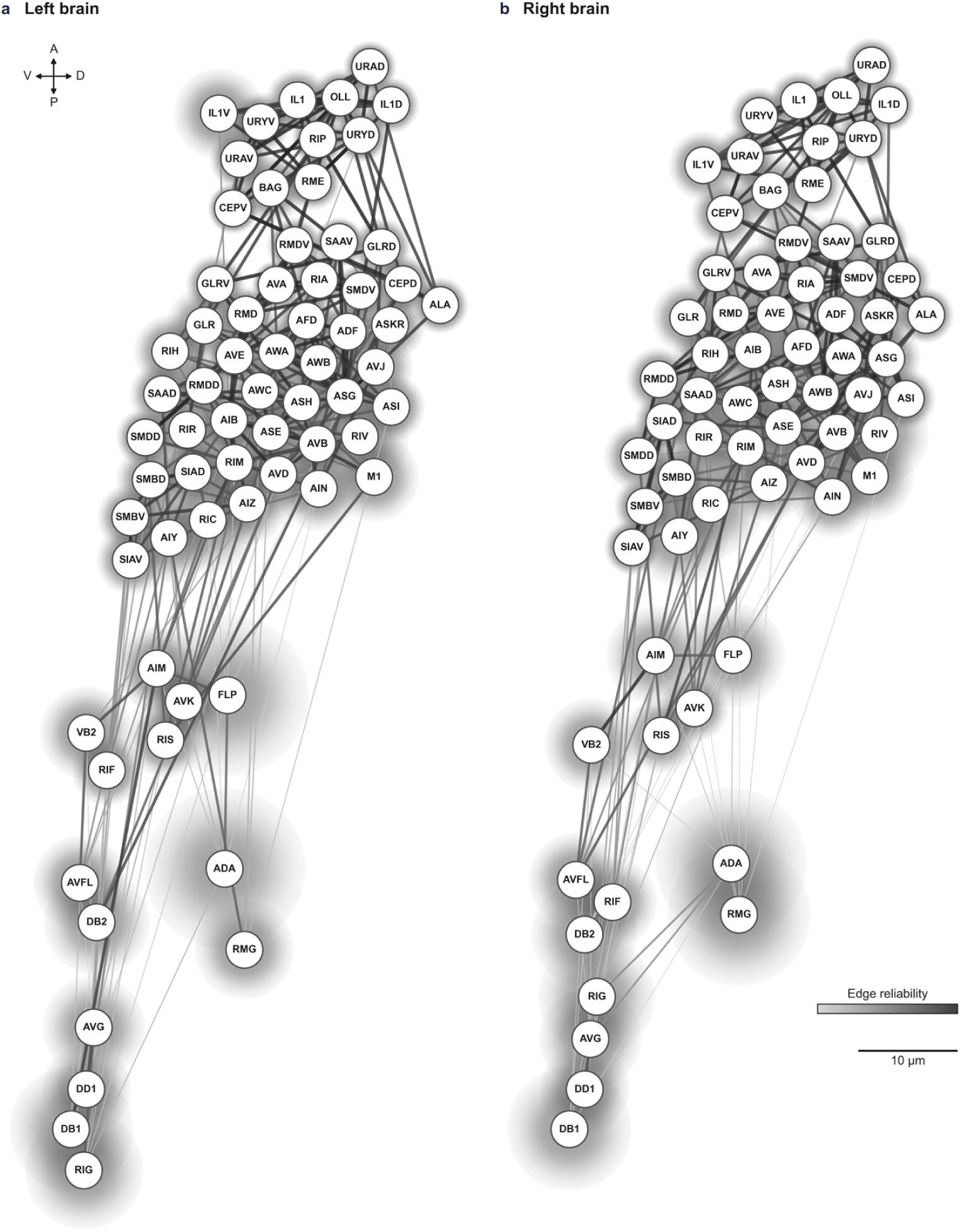
The brain-wide variability-aware atlas. Relational graph of the **a**. left and **b**. right neurons in the brain-wide variability-aware atlas.

